# Changing cognitive chimera states in human brain networks with age: Variations in cognitive integration and segregation

**DOI:** 10.1101/2025.04.28.650949

**Authors:** Andrew Patton, Jörn Davidsen

**Affiliations:** Complexity Science Group, Department of Physics and Astronomy, University of Calgary, Calgary, AB, Canada; Hotchkiss Brain Institute, University of Calgary, Calgary, AB, Canada

## Abstract

The aging process profoundly impacts the human brain, leading to notable changes in cognitive abilities. Although the brain’s structural and functional alterations with age are individually well documented, how differences in cognitive abilities emerge from variations in the underlying spatio-temporal patterns of regional brain activity is largely unknown. Patterns of increased synchronization between brain regions are taken as enhanced cognitive integration, while decreased synchronization is indicative of cognitive segregation. The ability to dynamically switch between different levels of integration and segregation across different cognitive systems is believed to be crucial for overall cognitive performance. Building on a recently proposed cognitively informed, synchronization-based framework, we study here age-related variations in dynamical flexibility between segregation and integration, as captured by changes in the variable patterns of partial synchronization or chimera states. Leveraging personalized brain network models based on large-scale, multisite datasets of cross-sectional healthy cohorts, we systematically show how regional brain stimulation produces distinct patterns of synchronization. We find that chimera states play a crucial role in regulating the balance between cognitive integration and segregation as the brain ages, providing new insights into the mechanisms underlying cognitive decline and preservation in aging. Whereas the emergent synchronization behavior of brain regions belonging to the same cognitive system often show the same aging trends, different cognitive systems can demonstrate distinct trends. This supports the idea that aging affects cognitive systems differently and that understanding this variability is essential for a more comprehensive view of neuro-cognitive aging. At the same time, dynamical flexibility increases in the oldest age groups across most cognitive systems. This may reflect compensatory mechanisms to counteract age-related cognitive declines and points towards a phenomenon of dedifferentiation. Yet, the multiplicity of behaviors highlights that whereas dedifferentiation emerges in certain cognitive systems, differentiation can also occur in others. This illustrates that these processes, though seemingly oppositional, can coexist and unfold in parallel across different cognitive systems.

**Author summary:** The healthy brain relies on a dynamic balance between two complementary processes—integration and segregation. Integration enables coordination between distributed brain regions, while segregation reflects specialization. Optimal cognitive function emerges from a highly flexible balance between these processes as implicated by greater cognitive performance. As we age, changes in cognitive abilities occur, especially at older age, while at the same time the brain reorganizes. The exact relationship between the two is largely unknown and our work aims to tackle this challenge. Using a data-informed computational model we examine age-related differences in dynamical flexibility between segregation and integration, as captured by changes in the variable patterns of partial synchronization called chimera states—patterns where some brain regions cooperate while others remain largely independent. Our research shows that chimera states are key to maintaining the brain’s balance between integration and segregation as we age. This balance shifts differently in brain systems responsible for distinct cognitive skills, which may provide new insights into the mechanisms underlying cognitive decline and preservation in aging. In particular, it supports the idea that aging affects brain systems differently and that understanding this variability is essential for a more comprehensive view of neuro-cognitive aging.

## Introduction

### An Aging Population

The global aging population is on a rapid climb, propelled by many factors: Declining fertility rates, advances in healthcare and medicine, and a maturing aging population entering the senior citizen threshold 65 years of age. Collectively, these factors contribute to this demographic shift [1]. Notably, the global population at large is experiencing a deceleration in growth, with approximately 10% already living in old age, a proportion anticipated to double by 2050. This demographic transition carries profound implications, as advancing age often brings an elevated vulnerability to health-related challenges, exerting significant strain on the well-being of seniors, caregivers, and the broader economy.

Addressing the multifaceted needs of an aging population necessitates robust infrastructure in social and health services [2, 3]. Successfully navigating this demographic shift—often referred to as successful aging [4, 5]—hinges on strategies implemented both early in life and in the current daily lives of older individuals. These strategies emphasize cumulative efforts, involving training and transitioning to a healthier, more social environment throughout the aging process to minimize decline [6]. Amid the myriad of concerns associated with aging, cognitive decline is particularly pressing, given its potential to accelerate debilitating conditions, such as Alzheimer’s [7, 8]. However, technological advancements are unveiling various mechanisms that the brain employs to protect itself from cognitive decline, boosting successful aging strategies.

### Cognitive performance and neurocognitive theories of aging

The brain can be considered as a complex network with a structural and dynamical topology that governs its functions; complex in the sense of its holistic, vast collection of 86 billion neurons [9] and a thousand to ten thousand times the connections between them [10]. These connections are organized in a very particular manner, enabling information traffic in the brain to flow efficiently [11]. Varying local connections to long-range connections across the brain [12], forming highly connected modules [13], and highly inter-connected hubs of neurons [14], bridging modules together, enable the dynamics of information flow. The brain operates within this modular, hierarchical organization through the interplay of two fundamental processes—integration and segregation. Integration, reflective of the interconnectivity between brain regions, facilitates the flow of information across both distant and local networks of the brain [15]. It plays a crucial role in coordinating diverse cognitive processes, ensuring stability in task execution through the consolidation of information [16]. This is especially important for managing tasks of greater complexity, such as working memory, and supports adaptive thinking by enabling cognitive flexibility [17–19]. In contrast, segregation refers to the organization of the brain into tightly connected communities of regions [15], each specializing in specific cognitive functions. Segregation is not only linked to cognitive specificity but also to processing speed and crystallized intelligence [20], and plays a key role in the efficient execution of specialized tasks, such as motor skills [16–18].

Dynamical segregation is most prominent during rest—when no specific task is being performed—and has been linked to more random, spontaneous activity, such as mind-wandering [16, 21]. Dynamical integration, on the other hand, occurs predominantly during a task, as the brain reconfigures itself, transitioning from a rest state to a task state. Once a task ends, the brain shifts from a more integrated state back to a more segregated state during rest [22]. Tasks require varying levels of integration and segregation, demanding high flexibility for successful performance [16, 23–26]. Indeed, to support diverse cognitive abilities, the brain needs to be able to reconfigure integration, segregation, and the balance between the two [20, 23]. At the same time, high dynamic flexibility is related to optimal creative thought and even positive mood [25, 26]. In the context of healthy aging, dynamic flexibility has been shown to be associated with cognitive performance: A stable form of flexibility is essential for optimal cognitive performance [27].

Yet, aging in the context of the dynamical interplay between segregation and integration, and the associated dynamical flexibility remains largely unexplored. How cognition is affected as the brain ages is instead usually examined through the realm of cognitive neuroscience, where three key theories hold significance in the context of successful advanced aging [28]: i) Reserves, encompassing both brain [29] and cognitive reserves [30]. Over the lifespan, the brain accrues beneficial quantitative measures, such as, increased white/grey matter volume, cortical thickness, and the density of neurons and synapses. Simultaneously, cognitive reserves, influenced by both genetic predispositions and environmental factors, accumulate throughout life, contributing to the creation of neural resources aimed at safeguarding cognitive function through compensation. ii) Maintenance [31, 32], a theory where genetic and environmental factors collaborate to shield neural reserves, preserving the youthful characteristics of a matured brain. iii) Compensation, representing the brain’s utilization of neural reserves to meet overwhelming cognitive demands [33–35]. These theories collaboratively construct a cohesive strategy for navigating the challenges of aging; however, the precise extent to which these underlying mechanisms operate with respect to dynamical flexibility in cognitive integration and segregation remains an open question.

### Cognitive Integration and Segregation as a Synchronization Phenomenon

Related to the question of aging in the context of the interplay between cognitive integration and segregation and the associated dynamical flexibility is how differences in cognitive abilities emerge from variations in the underlying spatio-temporal patterns of regional brain activity. Synchronisation, a phenomenon whereby neural assemblies—neurons on the smaller scale and brain regions on the larger scale—activate coherently in time [36], points towards their communication and collaborative effort, presenting a fingerprint of cognitive integration. Across the whole brain network, key to synchronisation is the large-scale integration of functionally specialized brain regions [37], the inter-connected modules needed to utilize flexible functionality. On the contrary, a lack of large-scale synchronisation is indicative of a highly segregated cognitive state, whose information flow is largely constrained to the local network for a specialized task [38]. For a functional, cognizant brain, integration and segregation must strike a balance—excessive integration (synchronization) can result in an epileptic seizure [39]; excessive segregation, on the other hand, does not enable flexibility necessary for a diverse set of cognitive tasks.

The notion of a chimera state, which is characterized by a co-existence of synchrony and asynchrony in a system [40–45], ties together this idea of dynamical flexibility between cognitive integration and segregation. Chimera states are states of partial synchronization arising from an underlying symmetry-breaking mechanism, which allows the system to dynamically select which units synchronize and which do not [46, 47]. This includes switching sets of oscillators between synchronized and asynchronized upon external stimulation [46], replicating phenomena of migratory bird, sea mammals, and human (the so-called “first-night” effect) unihemispheric sleep patterns [48–51]. This flexibility associated with chimera states can be directly related to the dynamical flexibility necessary to switch between different states of functional integration and segregation across cognitive systems [52]. In particular, a recently proposed cognitively informed, synchronization-based framework links chimera states and cognitive system functions that subserve human behavior by showing that all large-scale human cognitive systems can give rise to chimera states with various patterns of integration and segregation [52]. These patterns provide insight into how structural constraints within cognitive systems are linked to patterns of brain activity that reflect the systems’ ability to perform a variety of cognitive tasks.

Here, we build on this previous study [52] and investigate the aging brain through the same cognitively informed, synchronization-based framework. Specifically, we study age-related variations in dynamical flexibility between segregation and integration, as captured by changes in the variable patterns of chimera states. Leveraging personalized brain network models based on large-scale, multisite datasets of cross-sectional healthy cohorts, we systematically show how regional brain stimulation produces distinct synchronization patterns and how they change across age groups. The emergent synchronization patterns can be classified into three states–synchronous, asynchronous, and chimera—and we find that brain regions belonging to the same cognitive system often show the same aging trends in their synchronization behavior. Yet, different cognitive systems can demonstrate distinct aging trends in their synchronization behavior. At the same time, the synchronization behavior indicates a common shift toward more coordinated and stable brain activity as the brain begins its early process of maturing across all cognitive systems. We also find that the number (but not the proportions) of distinct synchronization patterns associated with chimera states is largely preserved for a given cognitive system across age groups, with the auditory and ventral-temporal cognitive systems exhibiting the highest degree of dynamical flexibility and the subcortical, default mode and attention cognitive systems exhibiting the highest degree of robust synchronization behavior. Yet, the dynamical flexibility exhibits age-related variations that are specific to a given cognitive system, despite an often observed trend of a significant increase in dynamical flexibility in the oldest age groups across many cognitive systems. This may reflect compensatory mechanisms to counteract age-related cognitive declines and points towards a phenomenon of dedifferentiation.

## Materials and methods

### Structural Connectomes

For our analysis, we take advantage of the open-source and standardized “10kInOneDay” database, which contains datasets of MRI neuroimages from 42 different research groups from around the globe [53]. All MRI neuroimages were processed using the same pipeline for diffusion weight imaging and T1 imaging, and the same methods were used to construct structural connectomes. The dataset in its entirety contains participants from across age groups, ordered in sequences of 5 years. We chose the datasets with the largest number of participants and with a high range across age groups, such that each age group had *>* 25 participants for that dataset. Consequently, in this study we utilize six datasets of healthy cross-sectional aging covering a wide range of ages from a minimum of 5 years to a maximum of 90 years. There are a total of *N* = 2018 structural connectomes used in this analysis. The age distribution of the participants is shown in Fig. 1.

**Fig 1.**
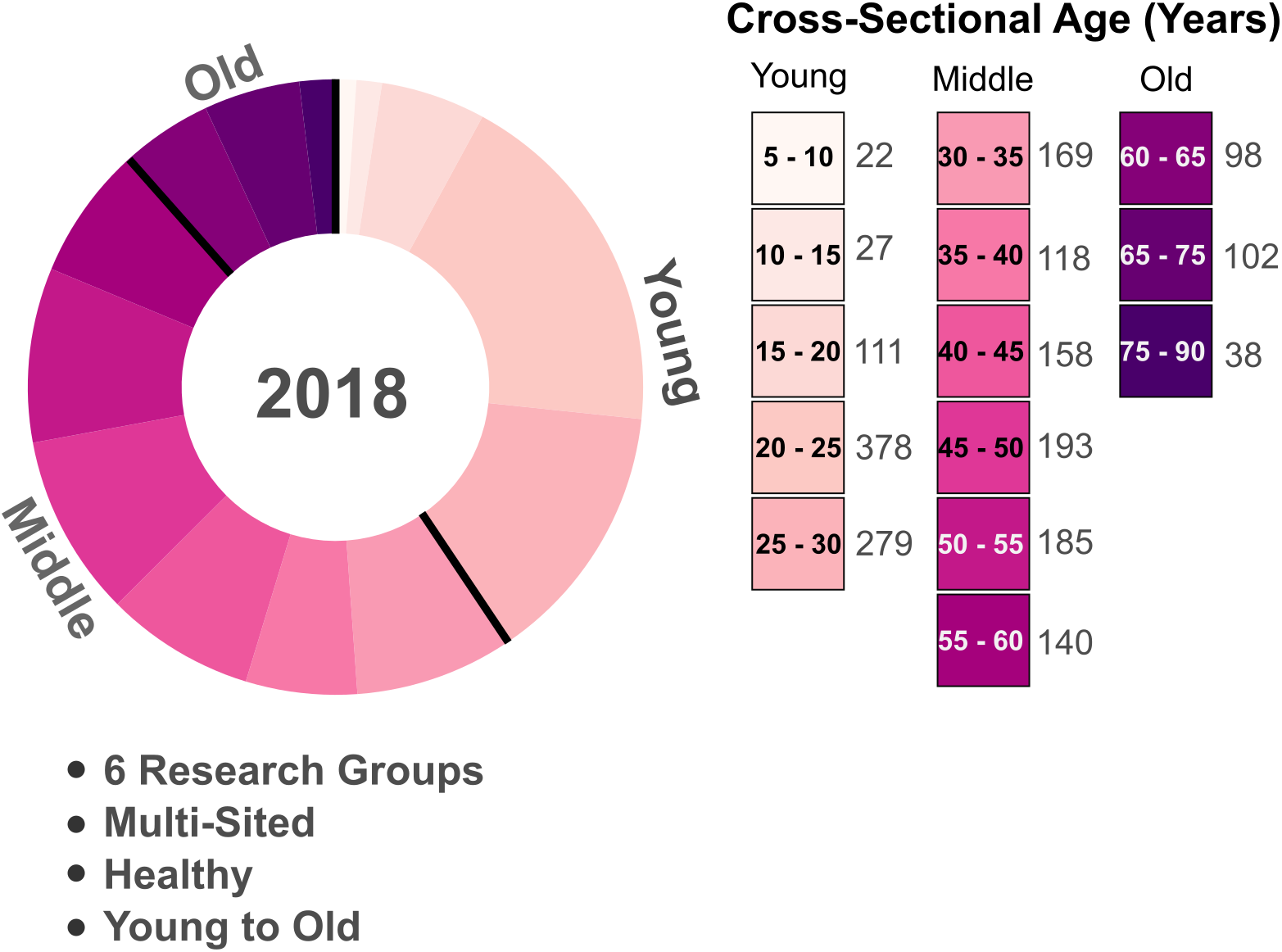
Cross-sectional age distribution of the ‘10kInOneDay’ dataset used in this study with additional details of the dataset. We consider a total of *N* = 2018 participants, which for some analyses we split to further categorize the ‘Young’ (*<* 30 years), ‘Middle’ (≥30 & *<* 60 years), and ‘Old’ (≥60 years) ages. Each participant is healthy. The dataset is multi-sited across 6 different research groups.

The structural connectomes contain 68 cortical regions from the Desikan-Killiany Atlas [54] and 14 subcortical regions from the subcortical Harvard-Oxford atlas. For a full list, please refer to Table SS1. The cortical regions were further fine grained from 68 to 114 using the Cammoun atlas [55] regions for a total 128 regions when including subcortical regions. Please note, when fine-graining, the same cortical regions are used as in the Desikan-Killiany atlas but the brain regions are further parcellated into sub-regions. For example, the right superior frontal brain region is parcellated into four sub-regions denoted by the subscripts ‘1’, ‘2’, ‘3’, and ‘4’. Each structural connectome consists of a network of brain regions (nodes) and the number of streamlines between each pair of brain region (undirected edges). To weigh the connections further, the number of streamlines connecting one brain region to another brain region is normalized by the sum of the two respective brain region volumes. Thus, each structural connectome is mathematically represented by an adjacency matrix *A*, where *A*_*jk*_ represents the weighted connectivity between brain region *j* and brain region *k*.

### Structural Connectivity and Rich Clubs

To quantify the topology of the structural connectomes and the cognitive systems, we can use different measures. One of them is strength which serves as a connectivity measure of a brain region. The strength of a brain region is defined as the total sum of its weighted connectivity, calculated using the following formula:

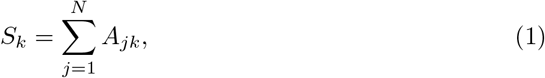

where *A*_*jk*_ represents the weighted connectivity between brain region *j* to brain region *k*.

The brain’s architectural organization is modular, characterized by densely interconnected clusters of brain regions forming a given module with sparse connections between regions in different modules [13]. Hence, for brain region *k*, its within-module or between-module connectivity can be quantified using the measure

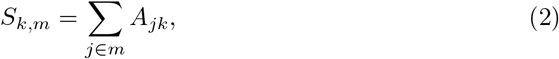

where *m* represents a specific module and the summation runs over all brain regions *j* within module *m*. Here, *S*_*k,m*_ measures the within-module strength of brain region *k* if *k* itself belongs to module *m*. Otherwise, if *k* is part of a different module, *S*_*k,m*_ instead quantifies the between-module strength from module *m* to the module containing brain region *k*.

A widely used measure for quantifying integration between modules is the participation coefficient. This metric measures the degree to which a brain region’s connections are distributed evenly among modules and is defined as

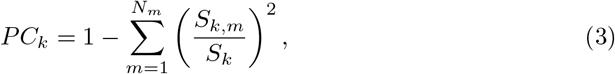

where *N*_*m*_ is the number of modules in a given structural connectome [56]. A high value indicates that node *k* acts as a connector between different modules, while a low value indicates that it is primarily connected to nodes within its own module.

Brain regions with a high strength that also act as the connectors of structurally segregated modules are considered hubs [57]. When a group of hub brain regions are more interconnected than is expected from their high connectedness, they form a so-called rich club [14]. In computational model studies, rich clubs have been reported to play a role in orchestrating synchronization of peripheral brain regions [58]. In particular, the shortest distance to the rich club is associated with the ability to induce synchronization where peripheral brain regions are less likely to synchronize upon a perturbation whereas perturbing brain regions close to the rich club core are more likely to facilitate global synchronization [52]. Thus, we also investigate the shortest distance to the rich club following the approach of determining the rich club of a structural connectome from previous studies [14, 59, 60].

Note while we considered other measures of the topology of the structural connectomes as well, such as betweenness centrality, local and nodal efficiency and the degree of nodes, none of these offered any additional insights beyond those arising from the strength distributions, the shortest distances to the rich club and the participation coefficient.

### Neural Stimulation in the Wilson-Cowan Neural Mass Model

The structural connectomes represent an in-situ, personalized brain that can be leveraged “in-silico” to simulate the nonlinear dynamics of large-scale brain activity. Such in-silico experiments can be employed to begin answering the question of how the brain responds to external stimuli [61] in a highly specified and controlled manner. Here, we pursue this avenue and closely follow previous work [52] using the Wilson-Cowan neural-mass model [62] to capture the mesoscopic nature of the structural connectome and brain regions in general. In this model, individual brain regions are governed by non-linear dynamics and coupled to other brain regions via the respective structural connectome. Given that each brain region contains a large number of both excitatory and inhibitory neurons, the phenomenologically derived model by Wilson and Cowan focuses on the average collective activity of these two neuronal populations and describes them by two ordinary differential equations with a sigmoid-shaped activation function,

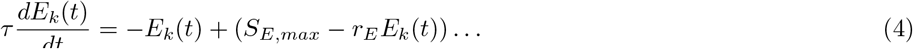

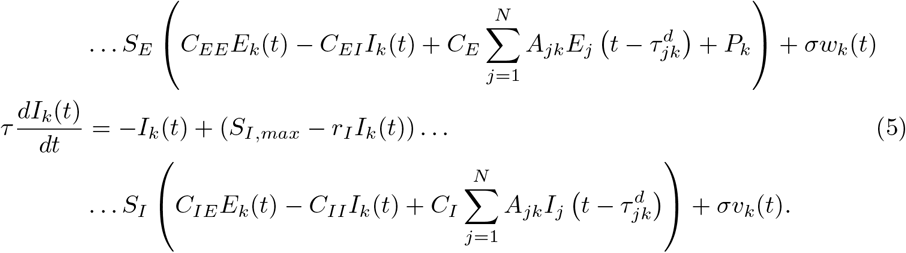

Here, *E*_*k*_(*t*) and *I*_*k*_(*t*) is the excitatory and inhibitory population activity for brain region *k* at time *t*, respectively. *τ* is a time constant, and *r*_*E*_ and *r*_*I*_ are the excitatory and inhibitory population refractory periods, respectively. *C*_*EE*_, *C*_*EI*_, *C*_*IE*_, and *C*_*II*_ are local parameters defining the self-coupling of the excitatory population, coupling between the excitatory population and the inhibitor population, coupling between the inhibitory population and the excitatory population, and the self-coupling of the inhibitory population, respectively, in a given region. Globally, the coupling strength between different brain regions is determined by *C*_*E*_ and *C*_*I*_. *A*_*jk*_ is the connection between brain region *k* and *j* from the structural connectome 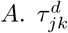 is the time delay between brain region *k* and *j* from the time delay matrix *τ*^*d*^, representing the time it takes for an electrical signal to travel between brain region *k* and *j* according to the conduction velocity of an unmyelinated axon (*v*_*d*_ = 10 ms^−1^ [63]). *P*_*k*_ is the stimulation strength for brain region *k*. Minimal white noise, *w*_*k*_(*t*) and *v*_*k*_(*t*), with small variance *σ* is added to the excitatory and inhibitory population, respectively, for spontaneous activity, representative of the resting-state brain. The sigmoid-shaped activation functions are given by,

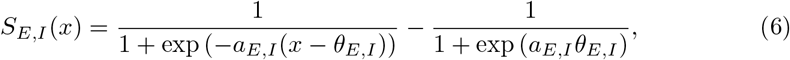

each representing the excitatory and inhibitory population, *S*_*E*_ and *S*_*I*_, respectively, with parameters controlling the mid-point and growth rate, *θ* and *a*, respectively, and

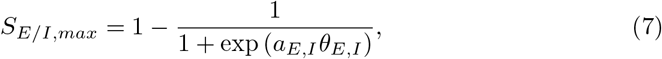

is the maximum value of the sigmoid for the excitatory and inhibitory population, respectively. Most of the parameters are biologically determined and here we use the same parameters as in reference [52]. The inhibitory global coupling term depends on the excitatory global coupling term, *C*_*E*_, such that *C*_*I*_ = *C*_*E*_*/*4, following that around four in five neurons are excitatory [60]. For each structural connectome, we select the value of *C*_*E*_ by tuning it to place the overall dynamics of the model in a “critical regime” using the midpoint method between the quiescent (the dynamics is at a fixed point) and active state (the dynamics is oscillatory) with a tolerance at 6 decimals points (see Fig. S1 for an example and the distribution of *C*_*E*_). Hence, in the absence of any external stimulation (*P*_*k*_ = 0 for all *k*) and without any internal noise (*σ* = 0) the overall dynamics remains at the fixed point corresponding to no activity across all brain regions (*E*_*k*_ = *I*_*k*_ = 0 for all *k*) but a sufficiently large stimulation of a single brain region will make that region oscillate. This means that a perturbation *P*_*k*_ of a single brain region *k* can cause a Hopf bifurcation, leading to limit cycle behavior of region *k*. If, however, the perturbation *P*_*k*_ is too large, the dynamics approach instead a new non-zero fixed point [64]. Therefore, we chose *P*_*k*_ = 1.15, which is sufficient to push the dynamics into an oscillatory regime. For each structural connectome, we investigate the emergent dynamics if a single brain region *k* is stimulated, i.e., *P*_*k*_ = 1.15 and *P*_*i*_ = 0 for all *i* not equal to *k*. This allows us to systematically stimulate each brain region individually and examine how the stimulation causes other brain regions to respond, as shown in Fig. 2 a) for an example. Some regions will also start to oscillate, while others do not and instead remain very close to the zero-activity fixed point.

**Fig 2.**
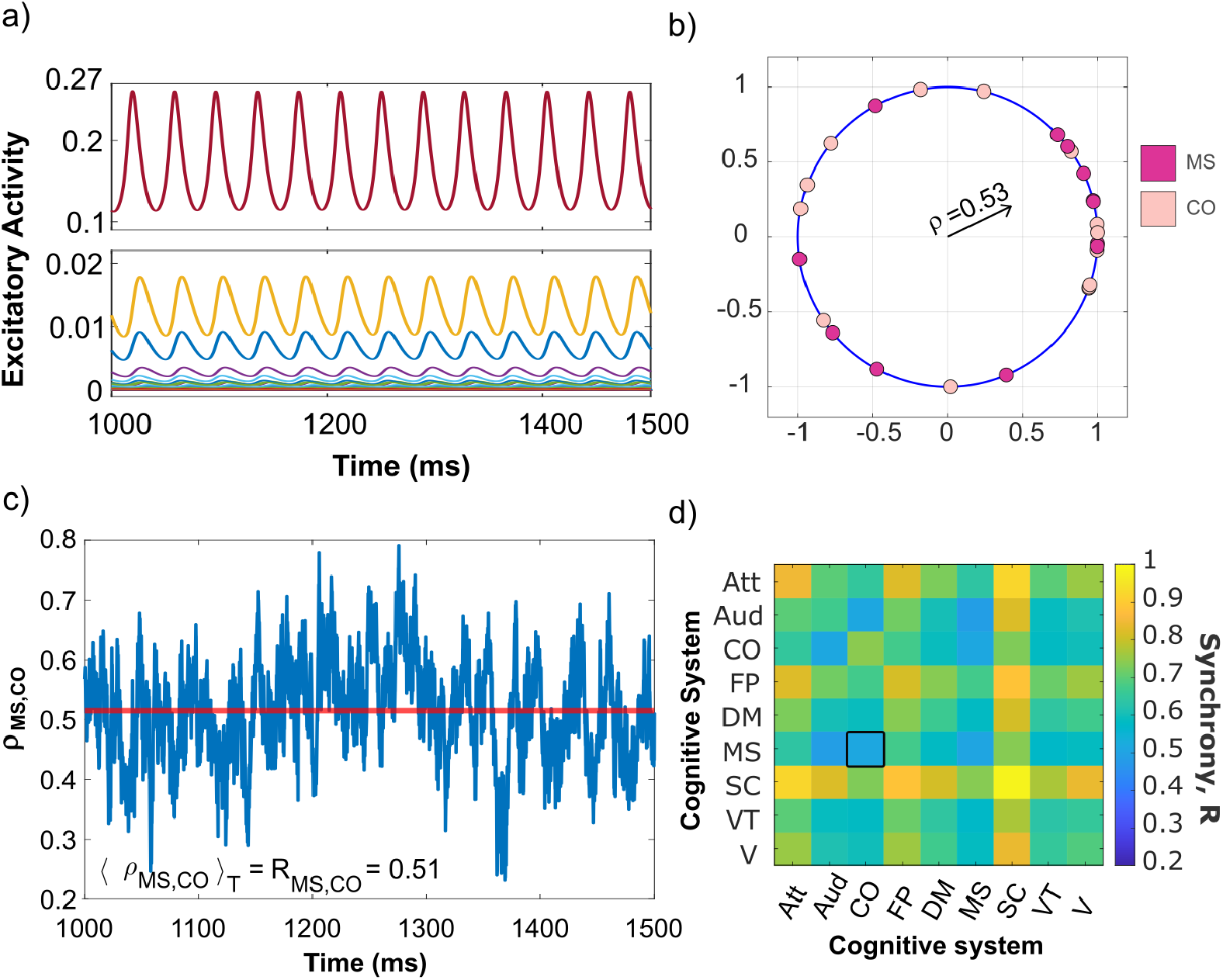
Synchronization framework: a) The excitatory time series for a single structural connectome after perturbing the left hemisphere insula subregion 1 brain region (belonging to the fronto-parietal cognitive system) with strength *P* = 1.15 for the entire duration. Top half shows the time series for the individually stimulated brain region and the bottom half shows the responding brain regions’ time series. Panel b) shows how to determine the amplitude of the complex Kuramoto order parameter, 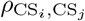, for a cognitive system pair, CS_*i*_, CS_*j*_. The brain regions that belong to the cingulo-opercular (CO) and motor-sensory (MS) cognitive system have a single snapshot in time of their oscillation phase (Eq. (8)) represented on the unit circle as dots, with CO in light pink and MS in dark pink. The length of the black vector arrow represents the amplitude of the complex Kuramoto order parameter, *ρ*_MS,CO_, measuring the synchronization with respective to the CO and MS cognitive systems. This example gives a value of *ρ* = 0.53 at for this snapshot in time. c) Time series of the amplitude of the complex Kuramoto order parameter, *ρ*_MS,CO_. The horizontal line in red, *ρ*⟨_MS,CO_⟩_*T*_ = 0.51, represents the mean over the time window, *T* = [1000, 1500]s, corresponding to the synchrony between the CO and MS cognitive systems. d) Matrix of the synchrony values for all pairwise combinations of the nine cognitive systems. The specific example for the synchrony between CO and MS is outlined with a black square.

For white noise (normally distributed), we want a minimal amount of variance, such that Euler-Maruyama integration remains applicable. Therefore, we set the variance *σ* = 1 × 10^−5^. As for simulation parameters, the time step is set at Δ*t* = 1 × 10^−3^ with initial conditions set at a small value of *E*_*k*_(0) = *I*_*k*_(0) = 0.1 for all *k* brain regions — away from its fixed point.

### Cognitive Systems Parcellation

For this study, we are interested in examining how the dynamics of the brain varies with age, and how this is organized with respect to cognitive function. To obtain such a cognitive picture, following a previous study [52], the 128 brain regions of the Cammoun atlas were grouped according to their well-documented functional connectivity that relates to their task-activated (or lack thereof) function [65, 66]. These so-called cognitive systems include the dorsal/ventral attention (Att), auditory (Aud), frontal parietal (FP), cingulo-opercular (CO), motor-sensory (MS), default mode (DM), visual (V), and ventral temporal (VT) systems. Since the subcortical regions are involved in this study for a more complete connectome, we grouped the subcortical regions into their own system called the subcortical (SC) system. Together, this gives nine cognitive systems that group all 128 brain regions, recorded in Table S1. Of note, these cognitive system assignments are kept fixed across age. We call these functional assignments cognitive systems to distinguish between the computational dynamics that are displayed here compared to the more empirical functional connectivity.

### Network Synchrony—Kuramoto Order Parameter

When a single brain region is stimulated, stable synchrony patterns can arise in response to this stimulation, such that multiple brain regions oscillate in a synchronized fashion. As reported in another study [52], the emergent patterns can be characterized as fully synchronous, fully asynchronous, or in-between these two extremes, termed chimera states. Using this framework, we systematically investigate how synchrony patterns change with age cross-sectionally.

To quantify the level of synchrony between brain regions and cognitive systems once a steady state has been reached, we use each brain regions’ steady-state time series, *E*_*k,ss*_(t) and *I*_*k,ss*_(t). To determine the phase associated with the possibly oscillatory dynamics of brain region *k* at time *t*, the geometric phase is utilized (the MATLAB function tan2), defined as

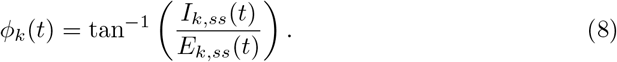

This allows us to determine how well synchronized the different cognitive systems are with respect to each other at a given time *t* through the so-called complex Kuramoto order parameter,

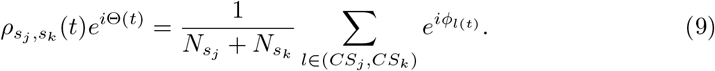

Here, *s*_*j*_ and *s*_*k*_ are different cognitive systems and *CS*_*j*_ and *CS*_*k*_ correspond to the sets of all brain regions that are part of *s*_*j*_ and *s*_*k*_, respectively. 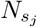 and 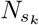 correspond to the number of brain regions in the respective set. Θ(*t*) is the average phase of those cognitive systems *j* and *k* at a time *t*. To obtain the amplitude, a measure of synchronization at time *t*, 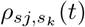, we simply apply the absolute value to Eq. 9,

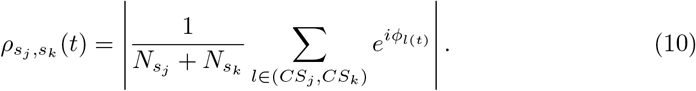

Using a time window of 500 ms, providing a sufficient number of oscillations (~13) to minimize the effect of fluctuations, we determine the total synchronization of the system, which we call the synchrony, through a time average of 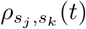,

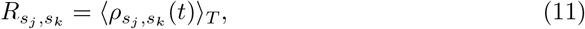

with *T* = 500ms, the length of the steady-state recording. By definition, 0 ≤ *R*≤1. Large values of the synchrony indicate a coherent or synchronous state, while small values indicate an incoherent or asynchronous state. For a visual example, consider 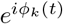,i.e., the dynamics of the phases on the unit circle, as shown in Fig. 2 b) for a single snap shot in time. If all brain regions oscillate with the same frequency and are phase-locked with zero phase lag, then their dynamics on the unit circle is identical and the system is in a completely coherent or synchronous state as indicated by *ρ*(*t*) = 1. Fig. 2 c) shows the time evolution of *ρ*_MS,CO_(*t*) and its corresponding synchrony for a single structural connectome as an example.

### Synchronization, Asynchronization, and Chimera States

When stimulating a single brain region and once the resulting synchrony measures are calculated between each pair of cognitive systems, a synchrony matrix can be constructed, as shown in Fig. 2 d). Even for the same structural connectome, the synchrony matrix can vary significantly depending on which brain region is stimulated, as shown in Fig. 3a), b), c) for three examples. To classify these emergent synchronization patterns and, hence, the stimulated brain region, we follow the same approach as [52]. First, synchrony matrices are binarized by applying a threshold; values of *R*≥ 0.65 are set to 1, representing a largely synchronous entry, while all other values are set to 0, representing largely asynchronous elements. The process of this transformation is illustrated in Fig. 3 where the pair-wise synchrony matrices in panels a, b, and c, are binarized producing the matrices in panels d, e, and f, respectively. All results reported in the following are qualitatively robust with respect to the value of the chosen threshold over an extended interval, [0.55, 0.80] (see also Fig. S2), confirming findings in a related study [52]. Using the binarized matrix, a generalized Louvain algorithm [67, 68] was applied to identify community assignments, as shown in Fig. 3g), h), and i). From these community assignments, cognitive systems may belong to the same community or a different community, capturing the synchronization patterns: Any cognitive system that belongs to a community with more than one member can be considered a member of the synchronous group ^1^, while any cognitive system that is grouped into its own community is a member of the asynchronous group. With the synchronous and asynchronous groups, cognitive patterns of synchronization are produced, as shown in Fig. 3j), k), and l). These synchronization patterns offer a representation of how the nine cognitive systems are (or are not) synchronized with one another. If the nine CS all belong to the asynchronous group, the individually stimulated brain region producing this pattern is classified as being asynchronous

**Fig 3.**
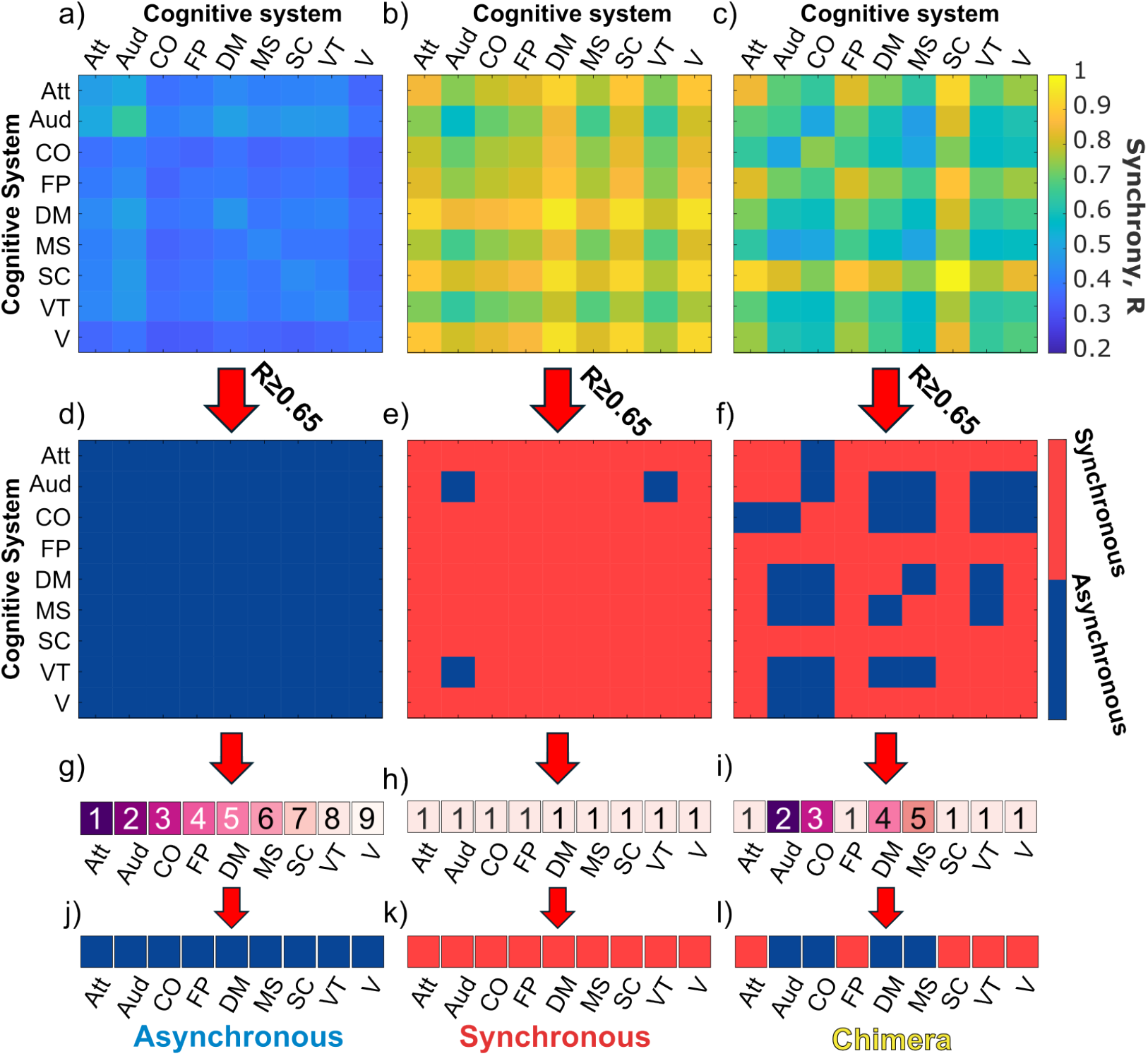
Classification of the emergent synchronization patterns when stimulating a single brain region for three different cases. The left column corresponds to stimulating the left hemisphere BANKSSTS, the middle columns is for the left hemisphere pallidum, and the right column is for the hemisphere insula subregion 1, all for the same structural connectome. Panels a, b, and c show their respective pair-wise synchrony matrices. Panels d, e, and f illustrate the binarization of the synchrony matrices using a threshold of *R* ≥0.65, with *R*≥ 0.65 representing a synchronous element (red) and an asynchronous element (blue), otherwise. Panels g, h, and i show the community assignments that occur when applying the Louvain algorithm. Cognitive systems belonging to the same community have the same colour and are labeled with the same number. Panels j, k, and l show the transformation of the cognitive systems’ community assignments into synchronization patterns, where blue colors indicate cognitive systems in the asynchronous group and red colors indicate cognitive systems in the synchronous group. The emergent synchronization patterns and, hence, the stimulated brain region are then classified as asynchronous (containing only an asynchronous group), synchronous (containing only a synchronous group), and chimera (containing both synchronous and asynchronous groups).

(Fig. 3j)). Conversely, if all nine CS belong to the synchronous group, the individually stimulated brain region is classified as synchronous (Fig. 3k). Any intermediate grouping, where there is a mix of synchronous and asynchronous group, will classify the individually stimulated brain region as a chimera state.

For each stimulated brain region a synchronization pattern is produced, and following this analysis pipeline, each of the 128 brain regions for all 2018 individuals are systematically stimulated to produce such synchronization patterns. Since each individual has a corresponding age, the structural connectome can be ordered according to the individual’s age. This allows us to investigate the cross-sectional differences of any given stimulated brain region and their emergent synchronization patterns. Here, we examine these age-related differences at different spatial scales—brain regions at the finer level and cognitive systems at the coarser level—and at finer and coarser age intervals. For the coarse-grained age intervals, the ages were aggregated into three groups: young, middle, and old, for any individual whose age is *<* 30 years old, ≥ 30 and *<* 60 years old, and ≥60 years old, respectively. At the finer-level, age intervals are incremented by 5 years, except for the older ages, whose ages are combined for more robust statistics (65 −70 & 70 −75 combine into 65− 75, and 75 −80, 80 −85, and 85 −90 combine into 75− 90), which in their 5 year bins alone would have been too low in sample size.

To capture the age-related differences in the classification as asynchronous, chimera, or synchronous of any given stimulated brain region, we focus on the proportion of each class in a given group, where the specific group depends on the chosen spatial scale and age intervals. Hence, the asynchronous, chimera, and synchronous proportion add up to 1 for a given age window and selected spatial scale.

From the abundance of synchronization patterns produced, we determine their prevalence by ranking the patterns for a given cognitive system and age window according to the number of times a pattern occurs and normalize it to obtain a proportion for easier comparison across cognitive systems. We can also determine the similarity between synchronization patterns across individuals and regions. Specifically, if in a given set of synchronization patterns a respective cognitive system always behaves in the same way (i.e., it is always synchronized or always asynchronized) then there is great similarity for that cognitive system. This can be extended to each of the nine cognitive systems to define the pattern similarity in a given set as follows:

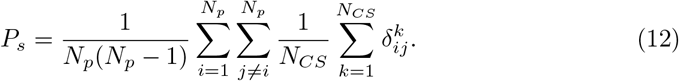

Here, *N*_*p*_ is the number of synchronization patterns in a given set and *N*_*CS*_ = 9 is the number of cognitive systems. 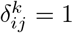 if cognitive system *k* behaves in the same way in pattern *i* and pattern *j*, otherwise 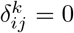. We will also consider the average pattern similarity across different sets, where each set can correspond to a different connectome (individual similarity) or to a different stimulated brain region (region similarity), for example.

### Potential Limitations

While the open-source “10kinOneDay” databse provided a substantial sample size (*N* = 2018), this strength also introduces a potential limitation: The aggregation of data from 42 international research groups using different scanners and field strengths introduces inter-group variability. To account for these scanner effects, we prioritized datasets with the large participant counts and broad age representation (ensuring *N >* 25 per age group), ultimately retaining 6 of the 42 datasets. Moreover, data harmonization methods—such as raw diffusion data correction or statistical adjustment for group differences [69, 70]—are commonly utilized to address such variability and we indeed found that scanner-related effects averaged out across the full cohort and did not significantly affect our results. Nevertheless, residual group-specific biases may be present.

Another limitation stems from the anonymization of participant ages into 5-year bins, which obscures finer-grained age-related trends. Future studies should validate our findings using datasets with continous age data. Additionally, the cross-sectional design limits causal inference or developmental findings, whereas a longitudinal analyses would help clarify whether observed patterns reflect true aging trajectories or cohort-specific effects.

A potentially critical limitation in developmental aging research, in particular, is selection bias in older cohorts due to survivor bias—individuals must survive to advanced ages to participate [71]. This introduces a confound where observed effects in older populations may reflect longevity-associated traits rather than standard aging processes, potentially distorting our age-related conclusions.

A further limitation arises from the scope of our computational model, which is exclusively focused on structural and neuronal properties. This framework does not include biological or chemical processes—such as metabolic changes, or molecular aging mechanisms—that are known to influence aging trajectories. Consequently, our findings may not fully generalize to empirical studies where these omitted factors could act as confounds. Future studies could integrate a multimodal computational model to bridge this gap but this currently presents a major challenge from both the modeling perspective and the data acquisition perspective.

## Results

We first examine the emergent synchronization patterns across brain regions and coarse-grained age windows using the pattern classification—asynchronous, chimera, and synchronous—described above. We find that while there is significant variability in the proportions and age-related changes in the proportions of the three classes between stimulated brain regions, those that belong to the same cognitive system typically share similar proportions, independent of the age window and despite changes in the proportions with age (Fig. 4). Subsequently, an analysis over more fine-grained age windows of selected cognitive systems exhibiting such consistent behavior highlights the differences and similarities in the age-related changes in the class proportion between the cognitive systems (Fig. 5). We investigate next how this consistent behavior relates to the topology of the underlying structural connectomes and their variations with age (Fig. 6, Fig. 7). Finally, we analyze which synchronization patterns typically arise for a given cognitive system and how they vary across age windows (Fig. 8). This allows us to answer the question whether cognitive systems are dominated by few patterns or rather many, sharing light on dynamic flexibility, as well as how the similarity of the synchronization patterns for a given cognitive system varies with age (Fig. 9). We further investigate to which degree these variations are present across regions within the same cognitive system and across individuals (Fig. 10).

**Fig 4.**
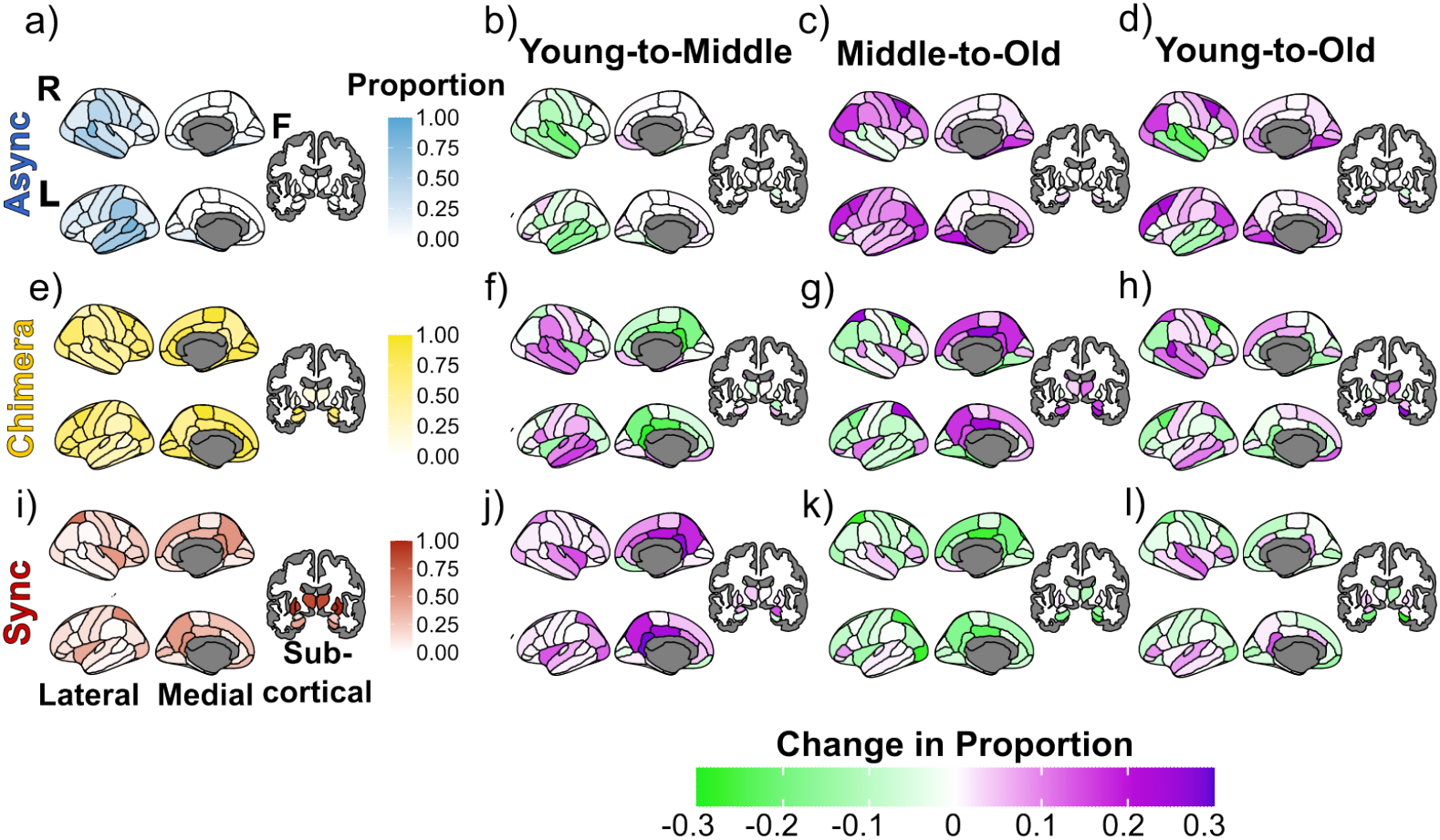
Proportions and age-related differences in the proportions of the three synchronization pattern classes associated with stimulated brain regions (coarse-grained atlas). The first column (panels a, d, and g) shows the average classification proportion with respect to the cross-sectional lifespan. The second column (panels b, e, and h) shows the change in classification proportion from young age (*<* 30 years) to middle age (≥30 & *<* 60 years) and the third column (panels c, f, and i) the change in classification proportion from middle age to old age (≥60 years), whereas the last column show the net change from young age to old age. The first row (panels a, b, and c) corresponds to the asynchronous classification proportions, the second row (panels d, e, and f) corresponds to chimera state classification proportions, and the third row (panels g, h, and i) corresponds to synchronous classification proportions. Each column contains three views for the brain: Lateral, medial, and subcortical, respectively from left to right. The lateral and medial view portray the right and left hemispheres, respectively from up to down, and the subcortical shows the front.

**Fig 5.**
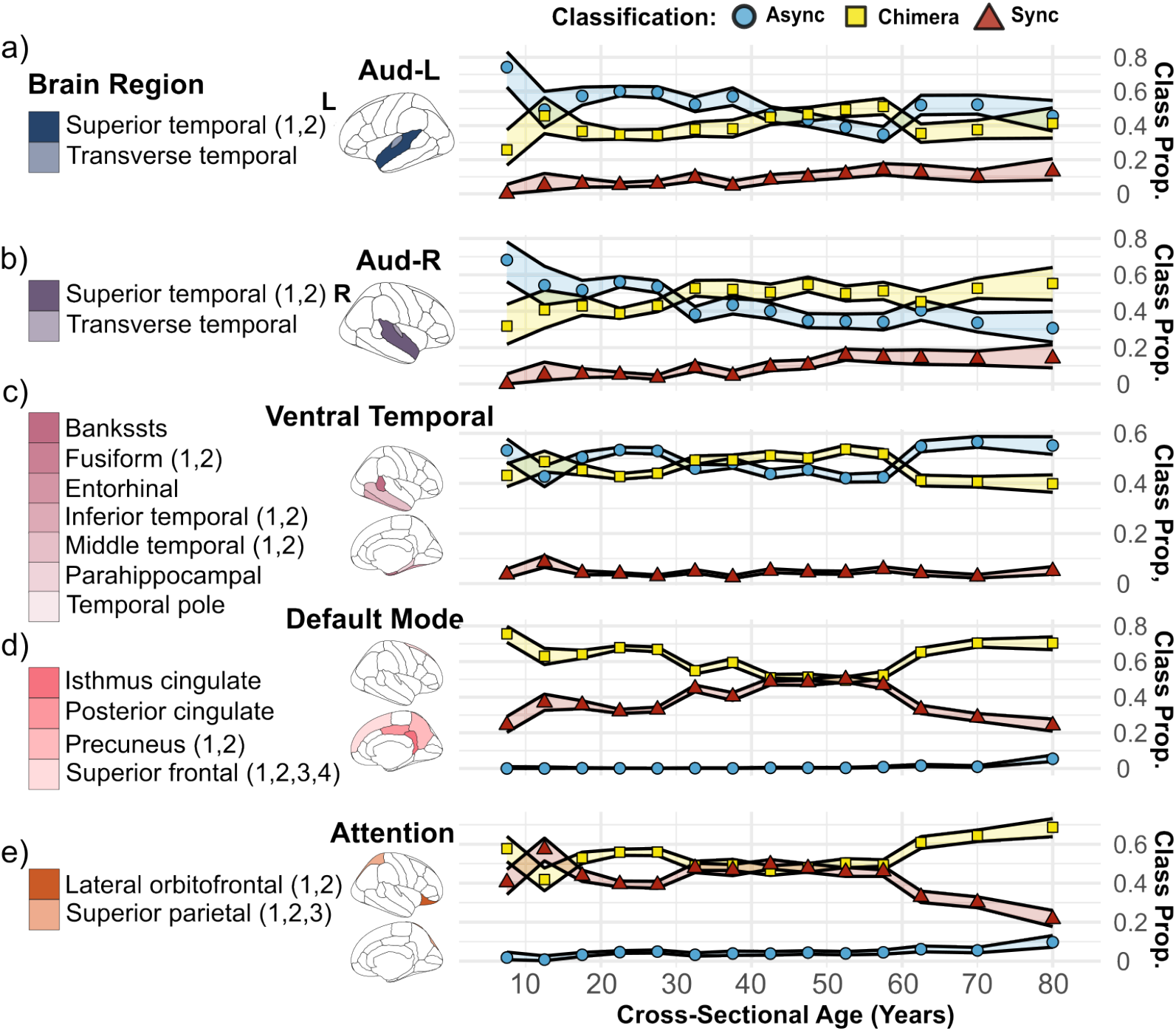
The classification proportions across cross-sectional age over five select cognitive systems: a) auditory-left (Aud-L; purple), b) auditory-right (Aud-R; blue), c) ventral temporal (magenta), d) default mode (pink), and e) attention (orange). Blue circles represent the asynchronous classification proportions, yellow squares represent the chimera state classification proportions, and red triangles represent the synchronous classification proportions. The first column shows the brain region labels that belong to the respective cognitive system for each cognitive system, the second column shows where the brain regions are located spatially, and the third column shows how the classification proportions change with cross-sectional age. Shaded regions are 95% confidence intervals across individuals.

**Fig 6.**
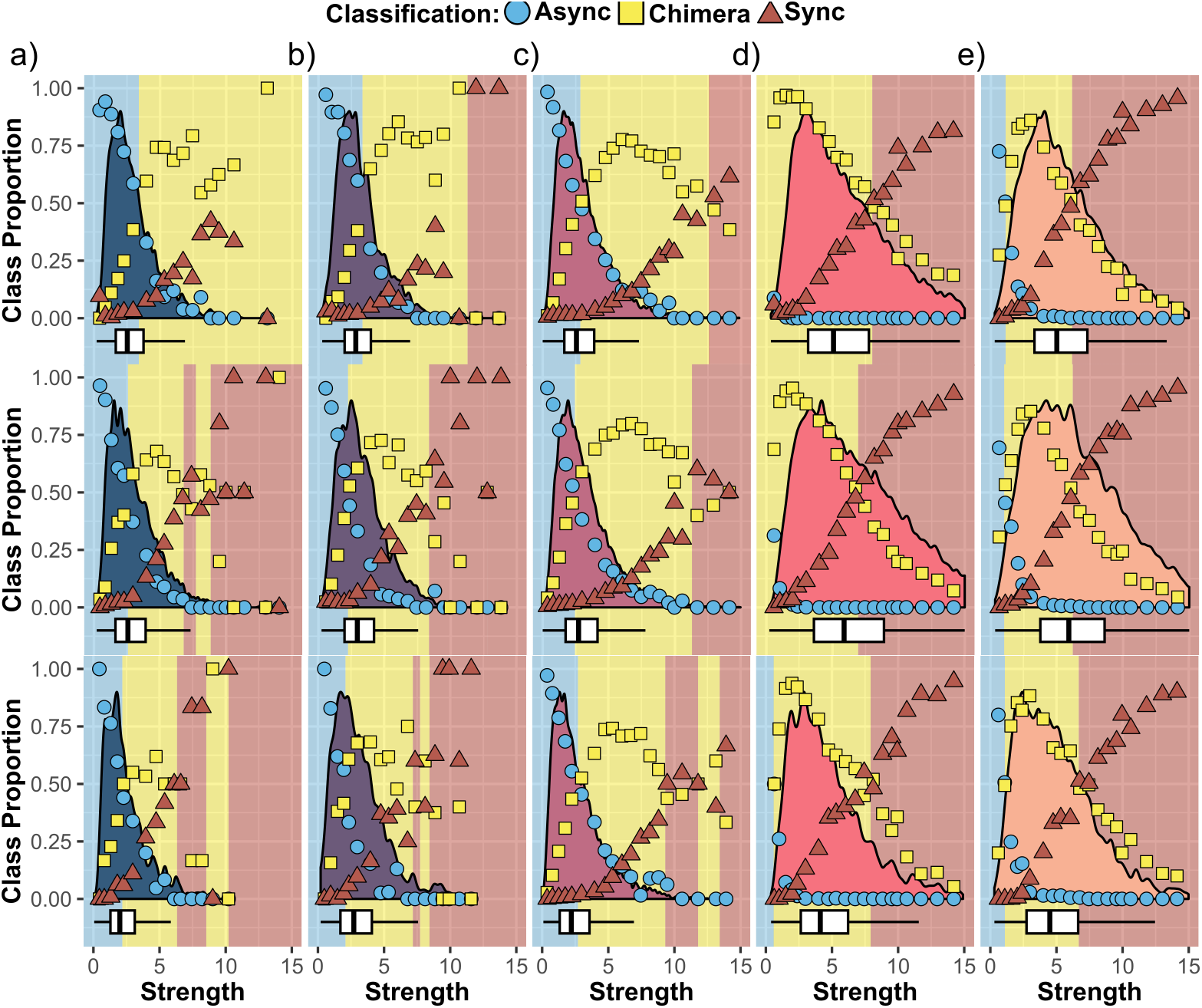
The variation in classification proportions with strength of the stimulated brain region belonging to the respective cognitive system in a given age group including the strength distribution along with its quartile as a box plot. Blue circles represent the asynchronous class, yellow squares represent the chimera class, and red triangles represent the synchronous class. Regions shaded in blue, yellow, and red are dominated by the asynchronous, chimera, and synchronous classification, respectively. Each column represents a different cognitive system a) for Auditory-Left (dark blue, b) for Auditory-Right (Dark purple), c) for Ventral-Temporal (Magenta), d) for Default Mode (Pink), and e) for Attention (orange). Each row corresponds to a different age group, from young (*<* 30 years) to middle-aged (≥ 30 % *<* 60 years) to old age (≥ 60 years).

**Fig 7.**
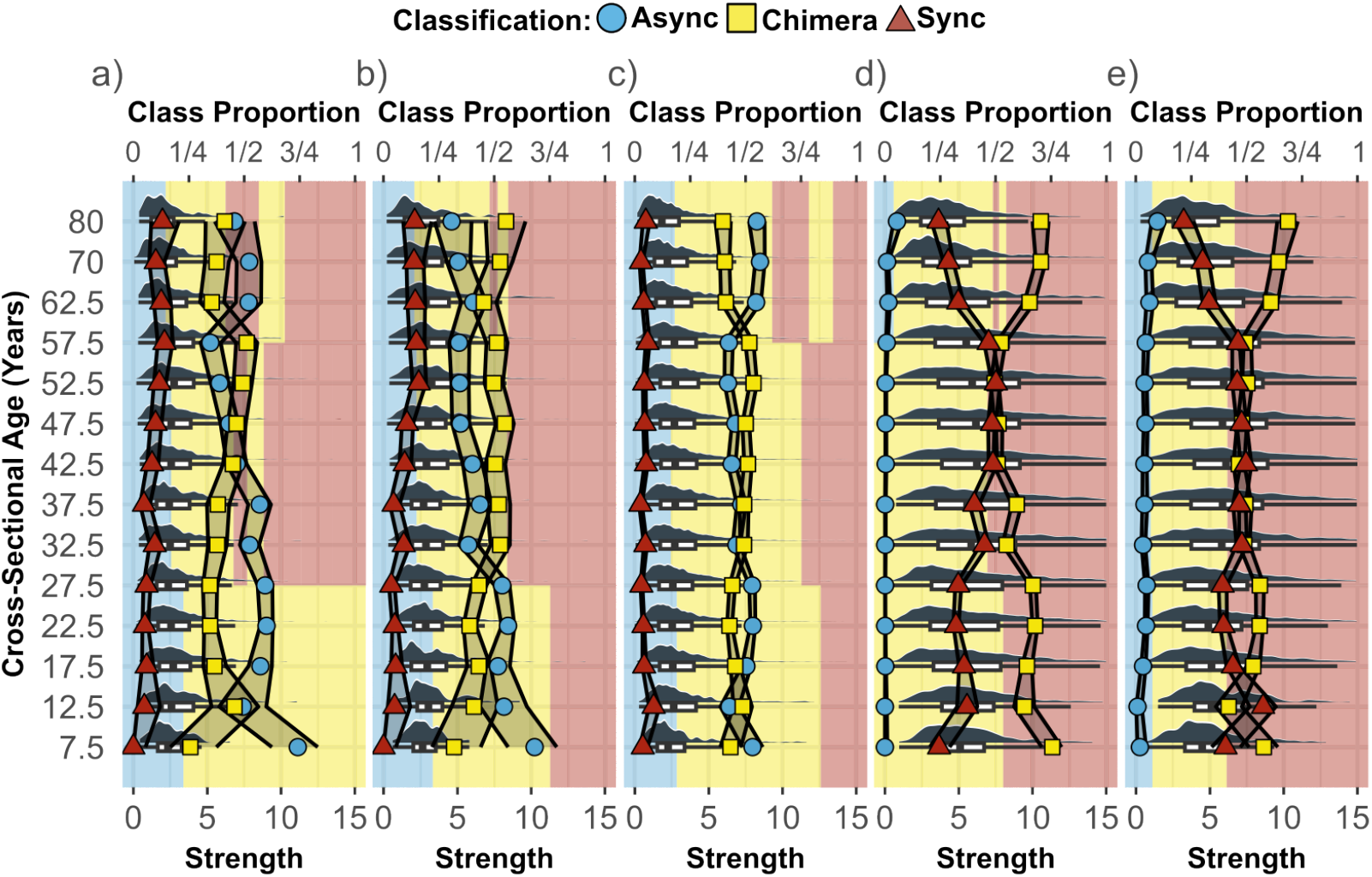
The variation in the dominating classification proportions with age and strength of the stimulated brain region belonging to the respective cognitive system including the classification proportions and strength distributions for a) the auditory-left, b) auditory-right, c) ventral-temporal, d) default mode, and e) attention cognitive systems. Shaded regions represent the dominating classification at that strength range (bottom x-axis) with blue, yellow, and red representing asynchronous, chimera, and synchronous classes, respectively. Symbols correspond to the classification proportions as in Fig. 5 (blue circles: asynchronous class, yellow squares: chimera class, red triangles: synchronous class, and top x-axis label). The strength distributions are shown (grey) along with their quartiles as a box plot.

**Fig 8.**
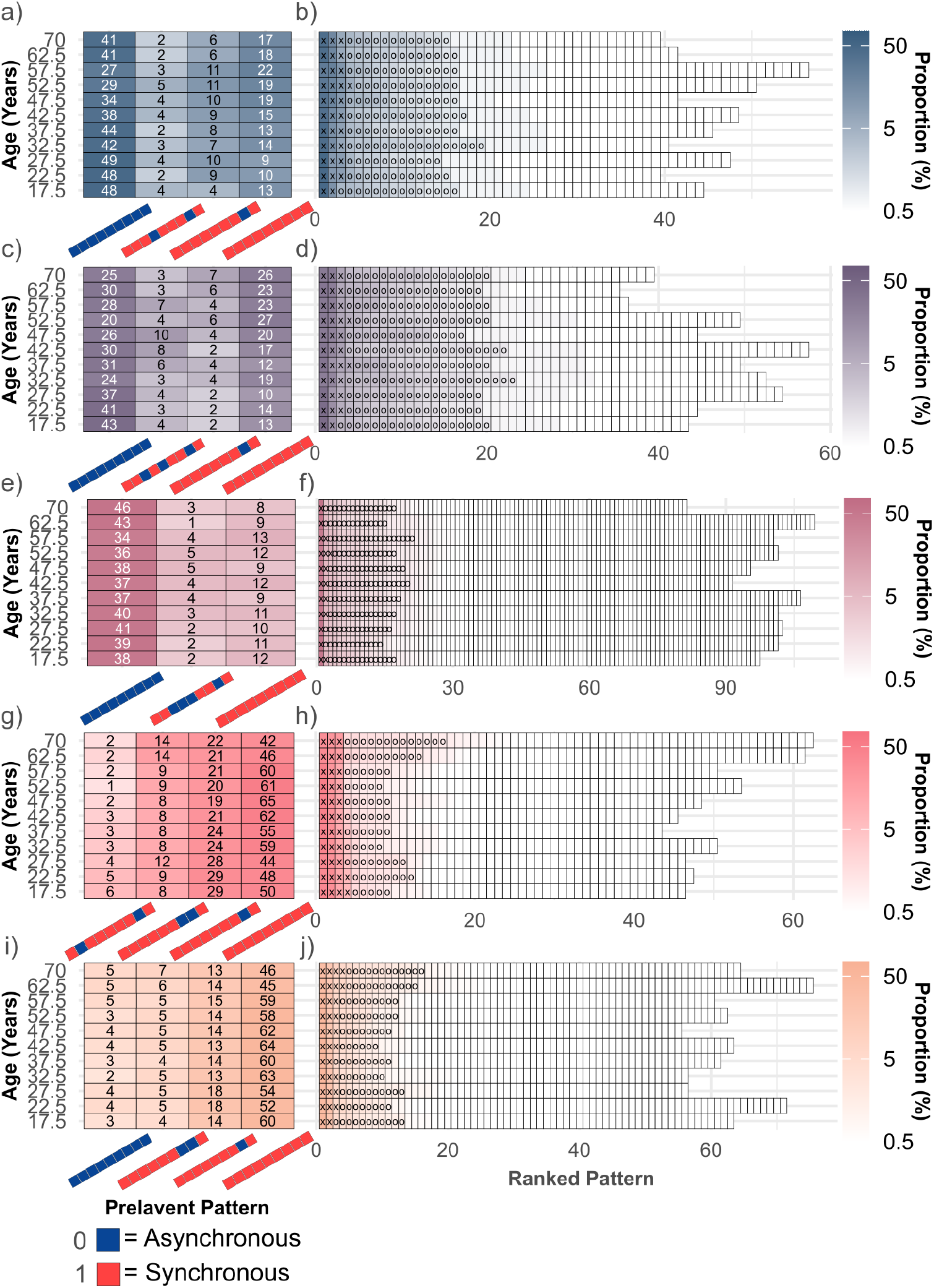
Proportion of emergent synchronization patterns along cross-sectional age (*N* ≈100 participants for each age group) for five different cognitive systems. The first column (panels a, c, e, h, i) shows the proportion of prevalent patterns (≥5% in at least one age window). As in Fig. 3, a synchronization pattern with a red (blue) block represents the synchronous (asynchronous) class. The second column (panels b, d, f, h, j) shows the proportion of patterns ordered by rank. Tiles containing an ‘X’ occur with ≥5% prevalence and ‘O’ occur with ≥3% prevalence. Each row represents a different cognitive system: auditory-left (panels a-b; dark blue), auditory-right (panels c-d; dark purple), ventral-temporal (panels e-f; dark magenta), default mode (panels g-h; pink), and attention (panels i-j; orange).

**Fig 9.**
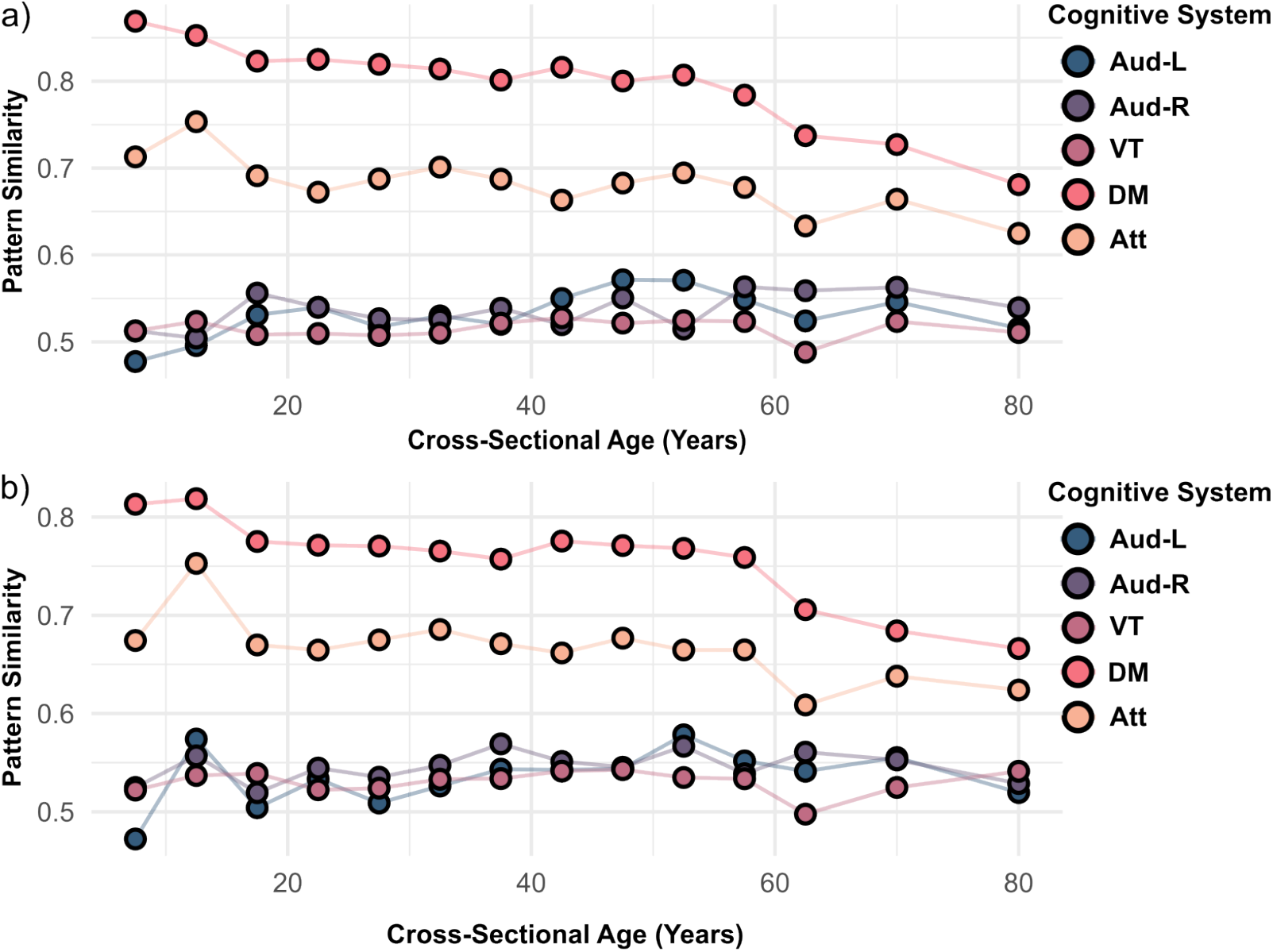
a) Variations in similarity of synchronization patterns, as defined in Eq. 12, with cross-sectional age across the Auditory-Left (dark blue), Auditory-Right (dark purple), Ventral-Temporal (magenta), Default Mode (pink), and Attention (orange) cognitive systems. b) Same as a) but for only the synchronization patterns classified as chimera.

**Fig 10.**
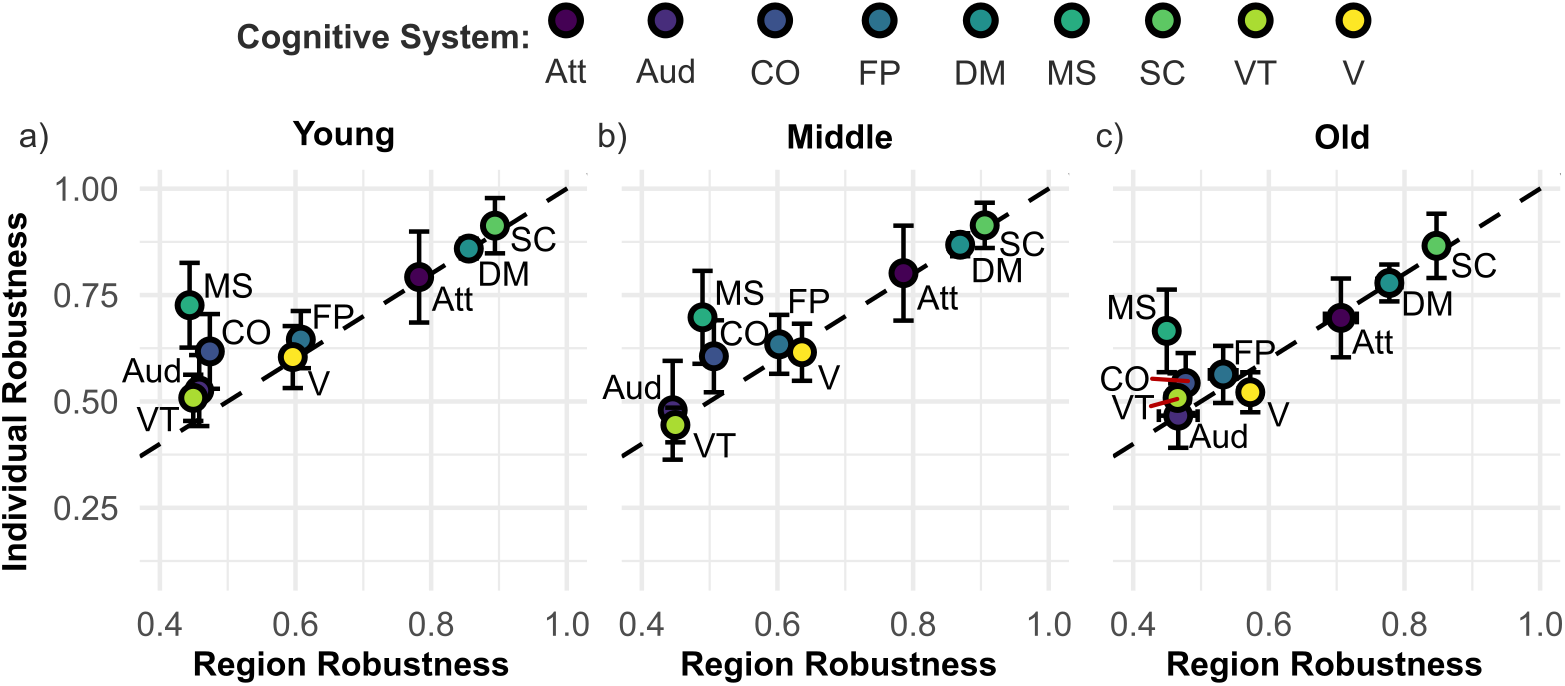
Region similarity and individual similarity across the three age groups: a) young (*<* 30 years, b) middle-aged (≥30 & *<* 60 years), and c) old age (≥60 years) for all cognitive systems: attention (Att), auditory (Aud), cingulo-opercular (CO), fronto-parietal (FP), default mode (DM); motor-sensory (MS), subcortical (SC), ventral-temporal (VT), and visual (V). Error bars show 95% confidence intervals. Note that the error bars in the x-direction are typically smaller than the symbol sizes.

### Age-related Changes in Synchronization Behavior Across Brain Regions

Focusing on the classification proportions associated with stimulated brain regions averaged across all ages first, Fig. 4a shows that asynchronous dynamics emerge most strongly when stimulating the temporal brain regions such as the banks of the superior temporal sulcus (bankssts), inferior-temporal, middle-temporal, superior-temporal, and transverse-temporal areas. These dynamics also appear, though to a lesser extent, in medial-ventral regions, including the parahippocampal, fusiform, and entorhinal cortices, consistent with the ventral-temporal cognitive system. Synchronous dynamics, by comparison, dominate when stimulating medial-top regions, such as the superior-parietal, precuneus, isthmus cingulate, posterior cingulate, superior frontal, and superior parietal cortices, aligned with the default mode and attention cognitive systems, respectively as shown in Fig. 4b. Strong synchronous activity is also evident when stimulating lateral regions like the insula and especially in subcortical structures, including the putamen, pallidum, thalamus, and caudate. Chimera states are broadly distributed throughout the brain and have particularly strong proportions when stimulating the hippocampus and amygdala structures compared to other subcortical regions as follows from Fig. 4c. Overall, while there is significant variability in the classification proportions between stimulated brain regions, those that belong to the same cognitive system typically share similar proportions.

Focusing on the age-related changes in the classification proportions of a given stimulated brain region next, Fig. 4 shows that brain regions belonging to the same cognitive system also demonstrate mostly consistent classification trends for asynchrony, synchrony, and chimera class proportions. The asynchronous proportion of brain regions exhibit complex, age-related differences, marked by distinct spatial and temporal patterns. As shown in Fig. 4b, from young (*<* 30 years) to middle age (≥30 and *<* 60 years), asynchronous proportions decrease overall, indicating a shift toward greater synchrony, manifesting itself through an increase in the proportions of the chimera class (Fig. 4f) and/or the synchronous class (Fig. 4j). This decline in asynchronous behavior is most pronounced in the superior temporal regions, for which the proportion of asynchronous behavior dominates when averaged across all ages as a comparison between Fig. 4a, e, i shows. However, from middle to old age (≥60 years), asynchronous proportions display a more nuanced trajectory (Fig. 4c), though there is an overall increase in asynchronous behavior largely driven by brain regions that had only small or no changes before. For superior temporal regions a hemispheric asymmetry emerges (at a significance level of *p <* 0.05): Asynchronous behavior in the upper temporal regions of the right hemisphere continues to decline, while that in the left hemisphere increases. For the inferior temporal, fusiform, parahippocampal, and entorhinal regions, the asynchronous proportion of brain regions additionally demonstrates notable age-related trends. The proportions follow nonlinear age trajectories, where asynchronous proportions initially remain relatively stable, declining weakly from young to middle age, and increase in older age. Asynchronous proportions also increase at older age in regions such as the lateral occipital, inferior parietal, caudal middle frontal, lingual, pericalcarine, and cuneus cortices. The net changes in the asynchronous proportions from young to old age are summarized in Fig. 4d. The bankssts, superior temporal, middle temporal, pars opercularis, and pars orbitalis brain regions show a decrease in their asynchronous proportions (as from young to middle age), while the rostral middle frontal, caudal middle frontal, inferior parietal, lateral occipital, lingual, inferior temporal, fusiform, parahippocampus, entorhinal brain regions increase in asynchronous proportions (as from middle to old age). From these brain regions, we see that changes in the asynchronous proportions arise in brain regions primarily in the auditory, ventral-temporal and visual cognitive systems.

The synchronous proportion of brain regions exhibits a clear age-related progression much like the asynchronous one. From young to middle age (Fig. 4j), synchronous proportions predominantly increase, which is especially pronounced in the default mode cognitive system’s brain regions including the isthmus cingulate, precuneus, posterior cingulate, and superior frontal, as well as in the lateral occipital, inferior parietal, and insula regions. However, there is a notable decrease in the medial orbitofrontal region during this period, marking an exception to the broader trend of increased synchrony. From middle to old age (Fig. 4k), however, a reversal is seen, with synchronous proportions decreasing in many of the regions that previously exhibited increases, such as the isthmus cingulate, posterior cingulate, precuneus, superior frontal, and inferior parietal regions. At the same time, synchronous proportions increase in specific areas, including the pars triangularis, pars orbitalis, and temporal regions. The age related changes from middle to old age are also characterized by a significant decline in synchronous behavior within subcortical regions, particularly the amygdala, thalamus proper, caudate, and hippocampus. The net changes in the synchronous proportions from young to old age are summarized in Fig. 4l indicating largely opposite changes to what is observed for the asynchronous proportions (Fig. 4d) but often with a lower absolute value.

The proportions of the chimera class (Fig. 4f,g,h) generally exhibit trends opposite to those of both the synchronous and asynchronous class, reflective of a compensatory relationship across brain regions, as chimera states are intermediary allowing synchronous and asynchronous dynamics in the brain to coexist. In brain regions where asynchronous proportions strongly decrease—such as the temporal regions—there is a corresponding increase in chimera state proportions, suggesting a shift in dynamics favoring hybrid activity patterns. Conversely, in areas where synchronous proportions strongly increase, such as the isthmus cingulate, chimera state proportions tend to decrease, indicating a reallocation of dynamics towards more global synchronization. Notably strong hemispheric asymmetries with opposite trends emerge in the changes with age in the lateral occipital region (at a significance level of *p <* 0.001). All these trends highlight the competitive balance among the three dynamical classes, where shifts in the asynchronous or synchronous class are often mirrored by compensatory changes in chimera states, underscoring their pivotal role in maintaining the system’s dynamical complexity across different brain regions.

### Age-related Changes in Synchronization Behavior Across Cognitive Systems

Given the aforementioned consistency of the age-related changes in the three class proportions for brain regions belonging to the same cognitive system for certain cognitive systems, we now focus on these systems including the auditory cognitive system though we split the latter into the left and right hemisphere due to the aforementioned asymmetry in the associated brain regions. In particular, we examine their age-related changes over more fine-grained age windows to allow for a more detailed examination of age-related changes in a given cognitive system. Fig. 5 shows this for the five selected cognitive systems (see Fig. S3 for the other cognitive systems). Notably, the auditory cognitive system, depicted in Fig. 5a for the left hemisphere and Fig. 5b for the right hemisphere, reveals significant hemispheric asymmetry in older age. Despite very similar and largely monotonic trends in the different classification proportions up to an age of about 55 for both hemispheres, the left hemisphere breaks with this trend and is largely dominated by asynchronous behavior at older ages, whereas the right hemisphere does not and is largely dominated by chimera behavior.

The ventral-temporal cognitive system, shown in Fig. 5c, follows a more nonlinear trajectory in its classification proportions. For example, asynchronous behavior declines in the youngest age bracket, stabilizes through adolescence, and peaks near the third decade, marking the start of the middle-age bracket. From that point onward, the proportion of asynchronous behavior gradually decreases until it begins to peak again in old age, around the sixth decade. The proportion of chimera behavior inversely mirrors these variations with age, while the proportion of synchronous behavior is small and largely constant. These non-linear variations reflect a dynamic and evolving balance of synchronization patterns within the ventral-temporal system throughout the lifespan.

A similar non-linear trajectory is observed in the attention cognitive system, see Fig. 5e, albeit the roles of the synchronous and asynchronous class are reversed. Here, a sharp increase in synchronous behavior is observed from the youngest age bracket through adolescence. From the start of the third decade until the sixth decade, the proportion of synchronous behavior remains largely constant until the sixth decade, where a pronounced decline occurs. The proportion of chimera behavior inversely mirrors these variations with age.

Finally, the default mode cognitive system, shown in Fig. 5d, is consistently dominated by chimera behavior and its proportion largely follows a distinct U-shaped trajectory. The proportion of synchronous behavior inversely mirrors this: Starting from a gradual incline that continues through the fourth decade, a plateau is reached in middle age, followed by a significant decline after the sixth decade.

Across all these five cognitive systems, a common trend at the youngest age groups (*<* 15 years) occurs. Namely, out of the two classes with the largest proportions the one that can be considered the more asynchronous one significantly decreases with increasing age, while the other one increases. Specifically, for the auditory-left, auditory-right and ventral temporal cognitive systems, the proportion of the asynchronous class decreases, while the proportion of the chimera class increases. Similarly, for the default mode and attention cognitive system, the proportion of the chimera class decreases, while the proportion of the synchronous class increases.

### Relating Age-related Changes in Synchronization Behavior to Changes in Structural Connectomes

To establish how the age-related changes in synchronization behavior observed above come about, we now investigate the influence of the topology of the underlying structural connectomes and their variation with age on the synchronization behavior. Specifically, we focus on measures that quantify the importance of a given brain region within the connectivity network of all brain regions captured by a given structural connectome. This allows us to relate the network properties of a brain region with the emergent synchronization patterns when that region is stimulated.

The relationship between classification proportions of the emergent synchronization patterns and the structural network strength of a stimulated brain region is shown in Fig. 6. In Fig. 6 a, b, and c, which depict the left hemisphere auditory cognitive system, right hemisphere auditory cognitive system, and ventral-temporal cognitive system, respectively, we observe that brain regions with low strength (low connectivity) are predominantly associated with emergent patterns that are asynchronous. In contrast, regions with very high strength (high connectivity) tend to lead to patterns that are classified as synchronous. Patterns classified as chimera states are most prevalent for stimulated brain regions with intermediate strength or connectivity, reflecting often a continuous transition from asynchronous to chimera and eventually synchronous behavior with connectivity strength in these cognitive systems. While this overall tendency is not surprising, note that all these cognitive systems exhibit substantial asynchronous behavior.

For cognitive systems that lead to predominantly synchronous activity with minimal asynchronous dynamics overall such as the default mode cognitive system (Fig. 5d) and the attention cognitive system (Fig. 5e), the associated brain regions tend to have larger strength values as intuitively expected, see Fig. 6d and e. At the same time, the range of strength values leading predominantly to chimera behavior or synchronous behavior is much broader. This indicates that the connectivity strength of a given brain region alone is not sufficient to determine which class of synchronization patterns will dominate when it is stimulated. Having in addition information about the cognitive system the brain region belongs to might also not be sufficient. This is because even within a given cognitive system, there are large age-related effects. Specifically, for the auditory and ventral-temporal cognitive systems (Fig. 6 a, b, and c), the range of strength values leading predominantly to asynchronous behavior tends to shrink with age, while the range dominated by synchronous behavior tends to expand. This can occur even in the absence of pronounced changes in the strength distribution with age (e.g., Fig. 6 b). In the attention cognitive system (Fig. 6 e), the synchronous region minimally decreases in size with age. Meanwhile, for the default mode cognitive system (Fig. 6 d), the synchronous region grows from young to middle age before shrinking again in old age. For both these cognitive systems, the strength distribution of the associated brain regions shows pronounced changes with age, with the oldest age group being dominated by smaller strength values. Fig. 7 investigates the relationship between strength distribution and class proportions further. In addition to the previously described trends, it also shows that even in the presence of strong age-related variations in the strength distribution and accompanied variations in the classification proportions the strength value ranges of the dominating class can remain largely unchanged (see, e.g., Fig. 7 e). Taking all these observations together, they suggest a dynamic redistribution of synchronous activity across the lifespan, influenced by regional connectivity and broader age-related changes in brain structure.

### Dynamical Flexibility: Emergent Synchronization Patterns and Age-related Changes

The synchronization patterns arising from stimulating a single brain region can vary significantly with the chosen brain region (see bottom row of Fig. 3). While we have categorized the patterns in three classes (asynchronous, synchronous, chimera), the number of synchronization patterns belonging to each class is not identical. Out of the total of 2^9^ unique patterns, two represent end members: One in which all nine cognitive systems are in the synchronous group and the other one in which all cognitive systems are in the asynchronous group. The former synchronization pattern is the only one classified as synchronous, whereas the latter patterns is the only one classified as asynchronous. The remaining 510 synchronization patterns are classified as chimeras, representing a range of different behaviors. This variety of chimera states in principle allows for dynamical flexibility between different levels of integration and segregation. Here, we analyze to which degree this is realized for different cognitive systems and how this varies across age windows.

We first focus on the contribution of different synchronization patterns, and based on their prevalence (Fig. S4) we categorize them into three groups: low-prevalence patterns (those contributing less than 3% to the total proportion), medium-prevalence patterns (3% – 5% contribution), and high-prevalence patterns (*>*5% contribution).

When many different synchronization patterns are present, this indicates high dynamical flexibility. Conversely, when a few high-prevalence patterns dominate, only few synchronization patterns account for most of the observed behavior, suggesting lower dynamical flexibility and a more stable and robust behavior. Hence, this approach allows us to characterize the degree of dynamical flexibility associated with chimera states and its variation with age, from enabling diverse neural strategies to robust behavior that supports consistent dynamics across individuals and/or brain regions.

Our findings are shown in Fig. 8 for five cognitive systems. Two striking observations emerge. First, the ventral-temporal cognitive system (Fig. 8 f) exhibits the greatest number of low-prevalence patterns, despite being dominated by the asynchronous pattern. This is then followed by the attention cognitive system (Fig. 8 j). In contrast, the auditory cognitive systems (left and right; Fig. 8 b and d, respectively) and the default mode cognitive system (Fig. 8 h) display noticeably fewer low-prevalence patterns. This trend is simply a consequence of the variable number of brain regions belonging to a given cognitive system. As indicated in Fig. 5, the number of regions is highest for the ventral-temporal cognitive system (10) and lowest for the auditory cognitive system (3). Focusing on the medium-prevalence and high-prevalence patterns instead, their number is largely age-independent and robust across cognitive systems and almost always lies in the range between 10 and 20. This is also true for the other cognitive systems (Fig. S5), with the auditory right and visual cognitive systems having the largest number of these patterns—suggesting a high degree of dynamical flexibility—whereas the subcortical cognitive system has the lowest number, suggesting more robust behavior.

The second observation in Fig. 8 pertains to high-prevalence patterns. A single chimera state pattern—characterized by an asynchronous ventral-temporal cognitive system while all other cognitive systems are in the synchronous group—prevalently appears across almost all cognitive systems. The only exception is when regions belonging to the ventral-temporal cognitive system itself are stimulated, which is not surprising given that brain regions within that cognitive system tend to behave very similar. The ventral-temporal cognitive system may be less responsive when other brain regions are stimulated due to being far from the rich club core of the brain and, more importantly, being more isolated compared to the other cognitive systems across all age groups as indicated by the lowest average participation coefficient (see Fig. S6). Beyond this, the synchronization patterns corresponding to the asynchronous and synchronous class are also among the high-proportion patterns for almost all cognitive systems. Only for the default mode cognitive system the synchronization pattern corresponding to the asynchronous class has a proportion of less than 1% across all age windows. Yet, for all cognitive systems and across all age groups the most prevalent pattern is one that belongs to either the synchronous class or the asynchronous class. For a given cognitive system this pattern is prevalent across all age groups, with the Aud-R being the sole exception where the synchronous pattern has a greater prevalence than the asynchronous pattern for age 52.5 only. High-prevalence patterns corresponding to the chimera class are rarer—there are at most two for each cognitive system in Fig. 8—and often exhibit variations in their prevalence with age though they all are characterized by an asynchronous response of the ventral-temporal cognitive system. Almost all of these observation are also true for the other cognitive systems (Fig. S5). This suggests that the dynamical flexibility associated with the synchronization patterns in the chimera class is often not static but exhibits age-related variations.

### Pattern Similarity

Even with a large number of distinct synchronization patterns, similarities between patterns can exist, reflecting either robustness through high similarity or variability through low similarity. Fig. 9 illustrates pattern similarity as defined in Eq. 12 across different ages for several cognitive systems (see Figs. S7 for the other cognitive systems). Notably, age-related changes in pattern similarity are evident for all cognitive systems. Overall, the shown cognitive systems can be grouped in roughly two sets, which align with the presence or absence of pronounced asynchronous class proportions (see Fig. 5, and Fig. S3 for the other cognitive systems). The auditory (left and right) and ventral-temporal cognitive systems are characterized by similarly low pattern similarity which remains largely constant with age when only synchronization patterns classified as chimera are considered. If all synchronization patterns are considered, slight differences emerge between them. The auditory (left and right) cognitive systems have a much higher pattern similarity at the youngest age, whereas the ventral-temporal cognitive system shows a systematic increase in pattern similarity starting around age 60. These differences are largely driven by the variations in the asynchronous class proportions of these cognitive systems with age as a comparison with Fig. 5 shows. In contrast, the attention and default mode cognitive systems are characterized by high pattern similarities, which remain largely constant up to age 60 before dropping to smaller values. This is independent of whether only synchronization patterns classified as chimera are considered or all patterns. As a comparison with Fig. 5d indicates, the variations in pattern similarity for those patterns classified as chimeras in the default mode cognitive system is not directly tied to variations in the associated chimera state proportions, which exhibits a distinct U-shaped trajectory with age such that the chimera state proportions are particularly high for the both the youngest age groups and the oldest ones. This observation indicates in particular that while the synchronization behavior is similarly dominated by chimera states in the oldest and youngest age groups in the default mode cognitive system, the set of chimera states is overall more diverse in the oldest age groups implying higher dynamical flexibility. It is also interesting to note that the degree of pattern similarity is not directly tied to the number of synchronization patterns observed for a given cognitive system as a comparison between the auditory (left and right) and ventral-temporal cognitive systems shows (see Fig. 8).

Low levels of pattern similarity for a given cognitive system can arise due to (i) significant variations in the synchronization patterns associated with stimulating different brain regions of that cognitive system for a given structural connectome, (ii) variations in the patterns associated with stimulating a given brain region across different structural connectomes, or (iii) a combination thereof. To examine this, Fig. 10 shows a scatter plot of the average pattern similarity across structural connectomes or individuals and across regions, respectively, for all cognitive systems and over different age groups. Most notably, the (average) region similarity and the (average) individual similarity for a given cognitive system take on largely identical values for almost all cognitive systems and across all age groups. This is indicative of case (iii), with both types of variations contributing equally. The only exceptions are the motor-sensory cognitive system, which consistently has a much higher individual similarity, and the cingulo-opercular cognitive system, which also has a higher individual similarity but becomes less pronounced with age. This implies that stimulating brain regions belonging to these two cognitive systems give rise to very distinct synchronization patterns for a given structural connectome but the emergent patterns are highly similar between different individuals (case (i)). Indeed, we find that for both these cognitive systems one subset of stimulated brain regions gives rise to patterns predominantly in the synchronous class, while another subset gives rise to patterns predominantly in the asynchronous class (Fig. S8). This features is highly reproducible across different individuals.

Fig. 10 shows that the similarity values are highest for the subcortical, default mode, and attention cognitive systems, which are characterized by an absence of pronounced asynchronous class proportions (see Fig. 5 and also Fig. S3). Their similarity values are statistically indistinguishable between the young and middle age group but decrease significantly in the old age group, consistent with the trend seen in Fig. 9 (see also Fig. S7). This indicates a loss of robustness and increase in variability for the old age group. The fronto-parietal and visual cognitive systems exhibit intermediate similarity values but different trends with age. While the fronto-parietal cognitive system remains constant before dropping for the old age group, the visual cognitive system exhibits a non-monotonic trend increasing first with age—especially in the individual similarity–before decreasing significantly in both region and individual similarity. This indicates a loss of robustness and increase in variability for the old age group for these two cognitive systems as well. The auditory and ventral-temporal cognitive systems consistently exhibit the lowest region similarity and remain constant across age groups within the given statistical uncertainties (for the separation into auditory-left and auditory-right cognitive systems, see Fig. S9). The motor-sensory and the cingulo-opercular cognitive systems exhibit intermediate individual similarity values but low region similarity values. Within the given statistical uncertainties the individual similarity remains constant with age, whereas the region similarity exhibits the same non-monotonic trend as the visual cognitive system. Since both these cognitive systems exhibit a dominating chimera class proportion of ≈60% across all age groups (see Fig. S3), the higher individual similarity indicates that the emergent synchronization patters in the chimera class (see Fig. S5) are largely consistent when stimulating the same given brain region across different structural connectome.

Comparing Fig. 10 with Figs. 5 and S3 indicates a relation between cognitive systems with a high synchronous class proportion (i.e., the default mode, attention and subcortical cognitive systems) tending to have high similarity values in their synchronization patterns, and cognitive systems with a high asynchronous class proportion (i.e., the auditory and ventral-temporal cognitive systems) tending to have low similarity values. All other cognitive systems have a comparable synchronous and asynchronous class proportion and tend to have intermediate similarity values. This suggests that a higher proportion of synchronous dynamics is indicative of lower dynamical flexibility, whereas a lower proportion indicates higher dynamical flexibility.

In summary, the age-related changes in pattern similarity reveal a distinct, cognitive system-specific behavior—some systems maintain a consistent, albeit low level of robustness, while others a more variable one. The former applies to the auditory and ventral-temporal cognitive systems. In older age, the default mode, attention, subcortical and fronto-parietal cognitive systems begin to lose some of their robustness both across regions and across individuals; this shift reflects a process of dedifferentiation, where these systems lose their distinctiveness and become less specialized in their dynamic patterns. At the same time, the visual and motor-sensory cognitive systems exhibit a non-monotonic trend, showing an increasing differentiation and greater distinctiveness in its patterns—especially when compared across brain regions in a given cognitive system—before decreasing again with age. This multiplicity of behaviors highlights an important insight: While dedifferentiation emerges in certain systems, differentiation can also occur in others—illustrating that these processes, though seemingly oppositional, can coexist and unfold in parallel across different cognitive systems.

## Discussion

Across the lifespan, in healthy aging, the brain experiences both structural and functional topological reorganization. Along side, there is an age-related progressive decline in cognitive performance. While there is a substantial body of literature on reorganization in functional topology [72–75] and to a lesser extent in the structural topology [76, 77], understanding the role of chimera states in cognitive aging and their varying manifestation in light of functional and structural age-related reorganization within cognitive systems and changes in dynamical flexibility remain limited. Here, by utilizing a large-scale, high-quality data set from a healthy cohort spanning ages from 5 to 90, we investigated for the first time age-related changes in both the emergent synchronization behavior under regional stimulation of personalized brain network models based on the Wilson-Cowan model and in the underlying structural connectomes of cognitively informed systems. Our aim was to establish a dynamical chimera state relationship within cognitive systems to explain age-related variations in dynamical flexibility between cognitive integration and segregation.

Our results reveal both universal trends—observed across virtually all cognitive systems—and system-specific effects that depend heavily on the cognitive system, pointing to inherent differences between them. Universally, regional brain stimulation reliably produces three distinct synchronization pattern classes: (i) complete synchronization, marked by globally coherent cognitive system activity; (ii) complete asynchronization, characterized by fragmented, independent dynamics; and (iii) chimera states, defined by the coexistence of synchronous and asynchronous clusters of cognitive system activity (Fig. 4a,e,i). These regimes map directly to principles of cognitive integration (via synchronization) and segregation (via asynchronization), with chimera states representing an intermediate regime that dynamically balances both processes—a hallmark of adaptive brain function. Across three distinct age groups (young vs. middle vs. old age in Fig. 4), regional cognitive integration and segregation largely follow a non-monotonic trajectory. Specifically, from young to middle age (Fig. 4 b,j) cognitive segregation declines as cognitive integration increases, while from middle to older age (Fig. 4 c,k) this relationship reverses—segregation rises and integration declines. These shifts are inversely mirrored by the reorganization of chimera states (Fig. 4 f,g), suggesting they play a role as a global transitional mechanism bridging fully synchronous pattern activity and fully asynchronous pattern activity through dynamic large-scale reconfiguration. Additionally, at the level of cognitive systems, a net increase in cognitive integration occurs across all cognitive systems from early childhood (5–10 years) to young adolescence (10–15 years; Fig. 5 & Fig. S3). This trend suggests a developmental shift toward heightened coordination and stability in brain activity, likely reflecting early neurodevelopmental maturation. These results are consistent with prior empirical reports of increasing integration during this developmental window [74].

Also at the level of cognitive systems, five cognitive systems demonstrate largely uniform dynamics across their associated brain regions and two distinct synchronization profiles emerge across these cognitive systems (Fig. 5): The auditory (separated into left and right hemispheres) and ventral-temporal systems exhibited high cognitive segregation—suggesting modular, domain-specific processing—whereas default mode and attention systems displayed high cognitive integration—reflecting broad cross-network coordination. These findings highlight how cognitive systems diverge in balancing segregated processing and integrative functions [74] and aligns with the default mode network’s reported role as a hub for global functional integration [78]. In old age, the auditory cognitive system exhibits a propensity for decreased cognitive segregation, favouring chimera states—a pattern that contrasts sharply with all other cognitive systems, all of which exhibit increasing cognitive segregation alongside a decline in cognitive integration, either through fully synchronous patterns or via chimera states (Fig. 5 & Fig. S3). This is consistent with a coexistence of differentiation and dedifferentiation as we will discuss further below.

### Hemispheric Asymmetries

In the auditory cognitive system, our results reveal age-dependent hemispheric asymmetries (Fig. 5): the left hemisphere exhibits a stronger propensity for cognitive segregation than the right, with this asymmetry amplifying across the lifespan and especially in old age. Strikingly, this increasing lateralization contrasts with the well-documented decrease in functional lateralization observed in many brain systems during aging—most notably the Hemispheric Asymmetry Reduction in Older Adults (HAROLD) phenomenon. HAROLD is characterized by reduced lateralization of prefrontal activity during cognitive tasks in older adults compared to younger individuals [79, 80], a pattern attributed to competing theoretical frameworks. Some propose compensatory mechanisms, such as the Compensation-Related Utilization of Neural Circuits Hypothesis (CRUNCH), where additional neural resources are recruited to maintain performance [81]. Others argue for age-related neural dedifferentiation or reduced processing efficiency [82]. Our findings, which highlight enhanced auditory asymmetry with age, diverge from these models. One possibility is that this discrepancy reflects limitations in capturing modulatory effects hypothesized to underlie age-related hyperactivation [83] or may not be neuronal in origin [84]. Testing this further presents an exciting challenge for the future.

### Age-related Differences in Structure

Our results indicate that the three synchronization pattern classes are linked to the underlying structure (Fig. 6). In particular, differences in the brain’s structural connectivity underpin the observed differences in synchronization pattern across different cognitive systems: Cognitive systems with low structural connectivity predominantly exhibit cognitive segregation (asynchronous dynamics), while regions with high connectivity favor cognitive integration (synchronous dynamics). Chimera states emerge in regions with intermediate connectivity, reflecting a continuous transitional regime that bridges the two other classes.

Our findings also reveal clear non-linear inverse U-shaped trajectories in strength with age—particularly prevalent in the default mode and attention cognitive systems (Fig. 7 & Fig. S6a). These non-linear patterns highlight periods of increased connectivity during midlife, followed by declines at older ages. Such trajectories align with previous reports of age-related U-shaped trends [60, 85]. On the other hand, the subcortical cognitive system follows a more monotonic decrease diverging from this non-monotonic behavior (Fig. S6a)). These results further emphasize the complex and dynamic nature of structural network organization across the lifespan—suggesting that specific cognitive systems may exhibit unique neuro-developmental trajectories in their structure. Notably, our results demonstrate that overall connectivity in the brain exhibits a pronounced decline with old age—a trend consistent with the well-documented effects of aging on structural networks and structural integrity [86–88]. While the overall structural connectivity contributes to cognitive integration and segregation trends, it alone cannot fully explain these trends. Instead, the cognitive segregation and integration emerges from a confluence of factors, including the topological architecture of the rich club—a core set of highly interconnected and thus influential brain regions—and its topological distance from the brain region being stimulated to the nearest brain region belonging to the rich club [52]. In our results, we largely observe a significant rise in older age for the distance to the rich club for each cognitive system (Fig. S6b)). This increase in rich-club distance suggests a weakening of the integrative hub functions that facilitate global communication, which may be a consequence of a dedifferentiation: As brain regions reorganize, the rich club, though containing intact hubs independent of age [60, 89], become more inefficient as the topological distance becomes greater, especially for the more core brain regions like those in the subcortical and default mode cognitive systems, resulting in differences of the cognitive dynamic flexibility. However, the properties related to the rich club can not explain differences in the cognitive systems with higher dynamical flexibility. For example, the ventral-temporal and visual cognitive systems are largely indistinguishable in their average connectivity strength and the average shortest distance to the rich club across all age groups but the average participation coefficient shows clear differences: The visual cognitive system acts more as a connector than the ventral-temporal cognitive system but loses this at old age, while the ventral-temporal system remains low in its participation coefficient value (Fig. S6). While all theses structural measures capture certain aspects related to dynamical flexibility they seem individually insufficient to explain cognitive integration and segregation.

### Synchronization-based Classification of Cognitive Systems

In terms of dynamical flexibility, our results, split into region robustness and individual robustness, show that while most cognitive systems have an individual robustness equal to the region robustness, there are four identifiable groups of interest (Fig. 10): One group exhibiting both large individual and region robustness (occupied by the subcortical, default mode, and attention cognitive systems), another group exhibiting both moderate individual and region robustness (fronto-parietal and visual cognitive systems), a group with both low individual and region robustness (auditory and ventral-temporal cognitive systems), and finally, off the diagonal, a group with low region robustness and moderate individual robustness (motor-sensory and cingulo-opercular cognitive systems). Notably, cognitive systems that belong to the first group produce predominantly synchronization patterns classified as synchronous (Fig. 8) and at the same time their synchronization patterns classified as chimera states are very similar (Fig. 9), whereas cognitive systems in the other groups produce a large number of synchronization patterns classified as asynchronous. Our results can be compared to a previous study [52], where, in terms of the regional and individual robustness of the synchronization patterns, the observations suggested the existence of potentially six groups, each one characterized as individually variable or individually stable based on individual similarity and characterized as cognitively diverse, cognitively flexible, or cognitively consistent based on region similarity. Of these potentially six groups, only four were observed [52]. Our results are consistent with the proposed grouping for the subcortical and attention cognitive systems (individually stable and cognitively consistent) and also for the cingulo-opercular and motor-sensory cognitive systems (individually variable and cognitively flexible) but not for the other cognitive systems. This might be due to the fact that the dataset in [52] only contained a small sample size of *N* = 30, compared to the lowest number of individuals of *N* = 238 in the smallest sample size of the age groups shown in Fig. 10. Moreover, our Fig. 10 explicitly shows the statistics with the mean and 95% confidence interval for both the region and individual robustness. Typically, the individual robustness is more variable and both confidence intervals get larger with smaller *N* such that the findings in [52] are consistent with ours if their statistical uncertainties are taken into account ^2^.

### Dedifferentiation and Differentiation in Cognitive Systems

Our investigation of changes in dynamical flexibility across age groups reveals a complex pattern of age-related reorganization across cognitive systems. Subcortical, default mode, and attention systems exhibit high functional specificity—reflecting their specialized roles—before slightly elevated cognitive variability occurs for the oldest age group (Fig. 10). In contrast, the ventral-temporal and auditory cognitive systems retain the same low functional specificity across all age groups. Notably, the default mode cognitive system’s ability of enduring pattern robustness from early adulthood into older age is likely critical for its role as a global integrator of cognitive processes [78]. However, at the oldest age group it is marked by a slightly diminished cognitive specificity (Fig. 10), increased chimera states (Fig. 5d), and heightened inter-individual variability, suggesting dedifferentiation—a loss of sharp distinctions between brain regions—and divergent dynamic trajectories that may reflect compensatory or degenerative pathways. On the other hand, the auditory cognitive system exhibits no significant age-related differences in its dynamical flexibility in both its region and individual robustness (Fig. 10) and the visual cognitive system exhibits an age-related decrease in its dynamical flexibility from young to middle age, indicating a process of differentiation, whereby the dynamics become more similar and thus distinctiveness increases across brain regions—suggesting a more specialized cognitive system.

Our findings on increased dynamical flexibility at older age are consistent with prior reports of age-related shifts in functional network topology, where youth is characterized by segregated networks that transition to integrated configurations in older age [74, 90, 91]. This reorganization is hypothesized to stem from neural dedifferentiation, wherein task-specific brain regions lose functional specificity and recruit broader networks [92–94]. For example, in a study on structure-function, an overcompensation effect is observed, wherein neural compensation leads to the activation of a broader array of task-unrelated cognitive systems, resulting in a network that is considerably less recognizable compared to its younger counterparts [95]. Such dedifferentiation parallels declines in cognitive aging, where reduced performance in cognitive abilities correlates with diminished neural specificity [96–98]. Our results extend this framework to the coexistence of both dedifferentiation and differentiation in different cognitive systems by linking dynamical flexibility and chimera states to the simultaneous decline and rise of cognitive specificity, suggesting that the coexistence of both of these processes may play a role in neurocognitive aging and thus may provide a more complete understanding to cognitive decline. Testing this directly remains an exciting challenge for the future.

## Conclusions & Key Takeaways

In conclusion, our main findings can be summarized in the form of two key points:

i. Existence of common and distinctive features in the trajectories of aging for cognitive systems: whereas brain regions belonging to the same cognitive system often show the same aging trends in their synchronization behaviour, different cognitive systems can demonstrate distinct aging trends in their synchronization behaviour. These findings support the idea that aging affects cognitive systems differently and that understanding this variability is essential for a more comprehensive view of neuro-cognitive aging. At the same time, the synchronization behavior indicates a common shift toward more coordinated and stable brain activity as the brain begins its early process of maturing across all cognitive systems.
ii. The role of chimera states in dynamical flexibility and aging: Chimera states provide a dynamical bridge between asynchrony and synchrony. Our findings indicate that the number (but not the proportions) of distinct synchronization patterns associated with chimera states is largely preserved for a given cognitive system across age groups, with the auditory and ventral-temporal cognitive systems exhibiting the highest degree of dynamical flexibility and the subcortical, default mode and attention cognitive systems exhibiting the highest degree of robust synchronization behavior. Yet, the dynamical flexibility exhibits age-related variations that are specific to a given cognitive system, despite an often observed trend of a loss of robustness in the oldest age groups across many cognitive systems. This may reflect compensatory mechanisms to counteract age-related cognitive declines and points towards a phenomenon of dedifferentiation, including in the default mode cognitive system, where cognitive integration and segregation are key. Yet, the multiplicity of behaviors highlights that whereas dedifferentiation emerges in certain cognitive systems, differentiation can also occur in others. This illustrates that these processes, though seemingly oppositional, can coexist and unfold in parallel across different cognitive systems.

## Acknowledgments

We would like to thank Andrea Protzner and Emma Towlson for helpful discussions. J.D. was supported by the Natural Sciences and Engineering Research Council of Canada (RGPIN/05221-2020). D.P. acknowledges financial support through an Alberta Graduate Excellence Scholarship and an Alberta Innovates Graduate Student Scholarship. The funders had no role in study design, data collection and analysis, decision to publish, or preparation of the manuscript.

## Data Availability Statement

The data used in this study are described in Ref. [53] and can be obtained upon request at dutchconnecomelab.nl. Specifically, the six datasets we analyzed from the “10kInOneDay” database are: 355581, 487062, 602004, 634143, 750313, 807122. The computational model used in this study is available in Ref. [52] in the data and materials availability section publicly available on Zenodo (doi:10.5281/zenodo.2590868; https://zenodo.org/record/2590869).

## Author Contributions

DP: Roles: Conceptualization, Formal analysis, Investigation, Methodology, Software, Validation, Visualization, Writing - original draft preparation, Writing - review & editing. JD: Roles: Conceptualization, Funding Acquisition, Methodology, Project Administration, Resources, Supervision, Validation, Writing - review & editing.

## Supporting information

**Table S1.** Cognitive system assignments for each brain region.

| Brain Region | Cognitive System |
| --- | --- |
| Lateral Orbitofrontal | Attention |
| Superior Parietal | Attention |
| Superior Temporal | Auditory |
| Transverse Temporal | Auditory |
| Caudal Anterior Cingulate | Cingulo-Opercular |
| Pars Opercularis | Cingulo-Opercular |
| Pars Orbitalis | Cingulo-Opercular |
| Rostral Anterior Cingulate | Cingulo-Opercular |
| Rostral Middle Frontal | Cingulo-Opercular |
| Supramarginal | Cingulo-Opercular |
| Caudal Middle Frontal | Fronto-Parietal |
| Inferior Parietal | Fronto-Parietal |
| Medial Orbitofrontal | Fronto-Parietal |
| Pars Triangularis | Fronto-Parietal |
| Frontal Pole | Fronto-Parietal |
| Insula | Fronto-Parietal |
| Isthmus Cingulate | Default Mode |
| Posterior Cingulate | Default Mode |
| Precuneus | Default Mode |
| Superior Frontal | Default Mode |
| Paracentral | Motor-Sensory |
| Postcentral | Motor-Sensory |
| Precentral | Motor-Sensory |
| Thalamus | Subcortical |
| Caudate | Subcortical |
| Putamen | Subcortical |
| Pallidum | Subcortical |
| Hippocampus | Subcortical |
| Amygdala | Subcortical |
| Nucleus Accumbens | Subcortical |
| BankSSTS | Ventral-Temporal |
| Entorhinal | Ventral-Temporal |
| Fusiform | Ventral-Temporal |
| Inferior Temporal | Ventral-Temporal |
| Middle Temporal | Ventral-Temporal |
| Parahippocampal | Ventral-Temporal |
| Temporal Pole | Ventral-Temporal |
| Cuneus | Visual |
| Lateral Occipital | Visual |
| Lingual | Visual |
| Pericalcarine | Visual |

These cognitive system assignments correspond to both the left and right hemispheres of the brain, as well as the fine-grained atlas, containing the same brain region name. For example, the brain region 'L insula 1' and 'L insula 2' belong to the brain region 'insula' in this table and thus to same cognitive system.

**Fig S1.**
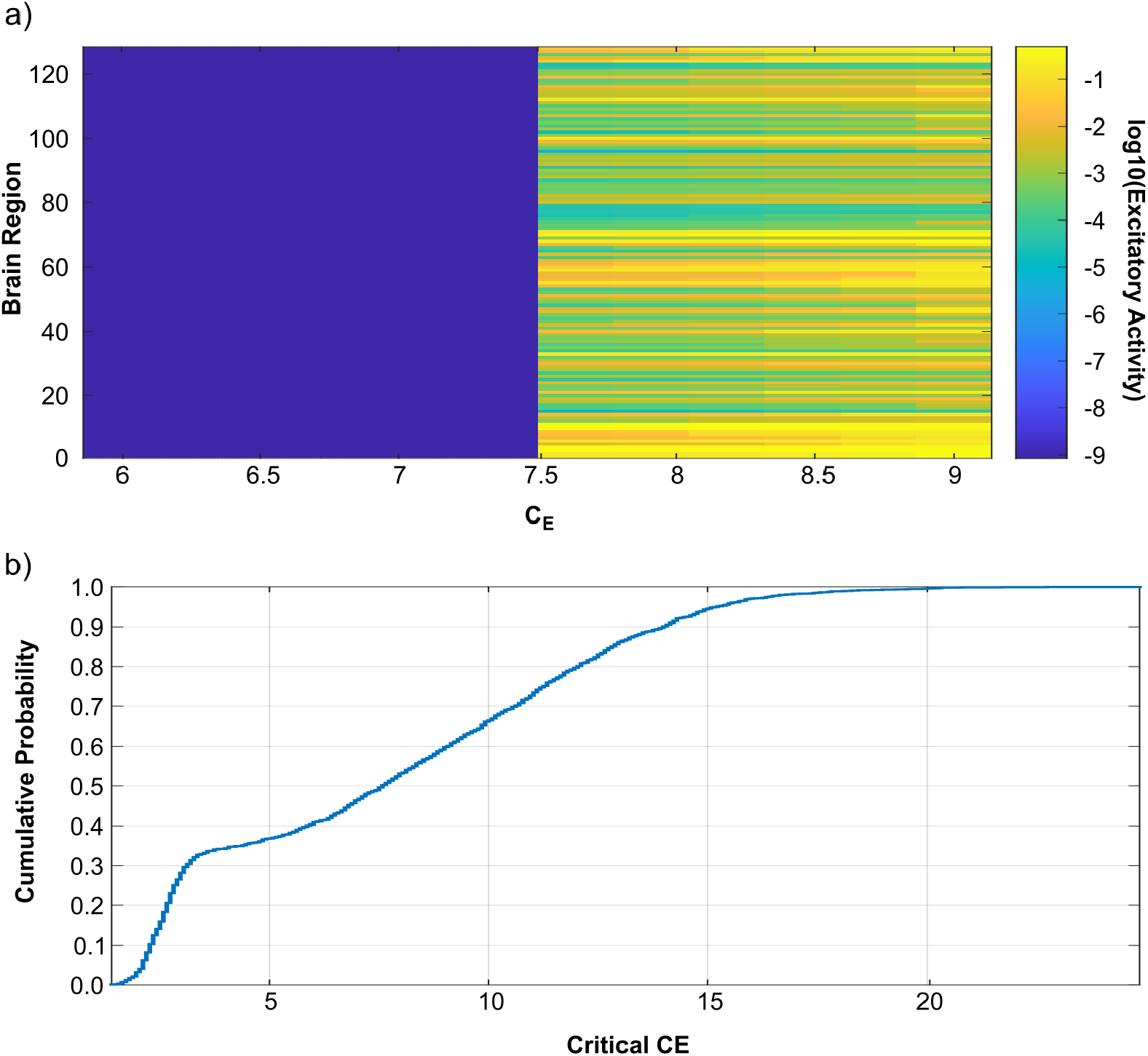
a) Example of excitatory activity for each brain region as a function of *C*_*E*_. b) Cumulative distribution function of the critical values of *C*_*E*_ for all *N* = 2018 structural connectomes.

**Fig S2.**
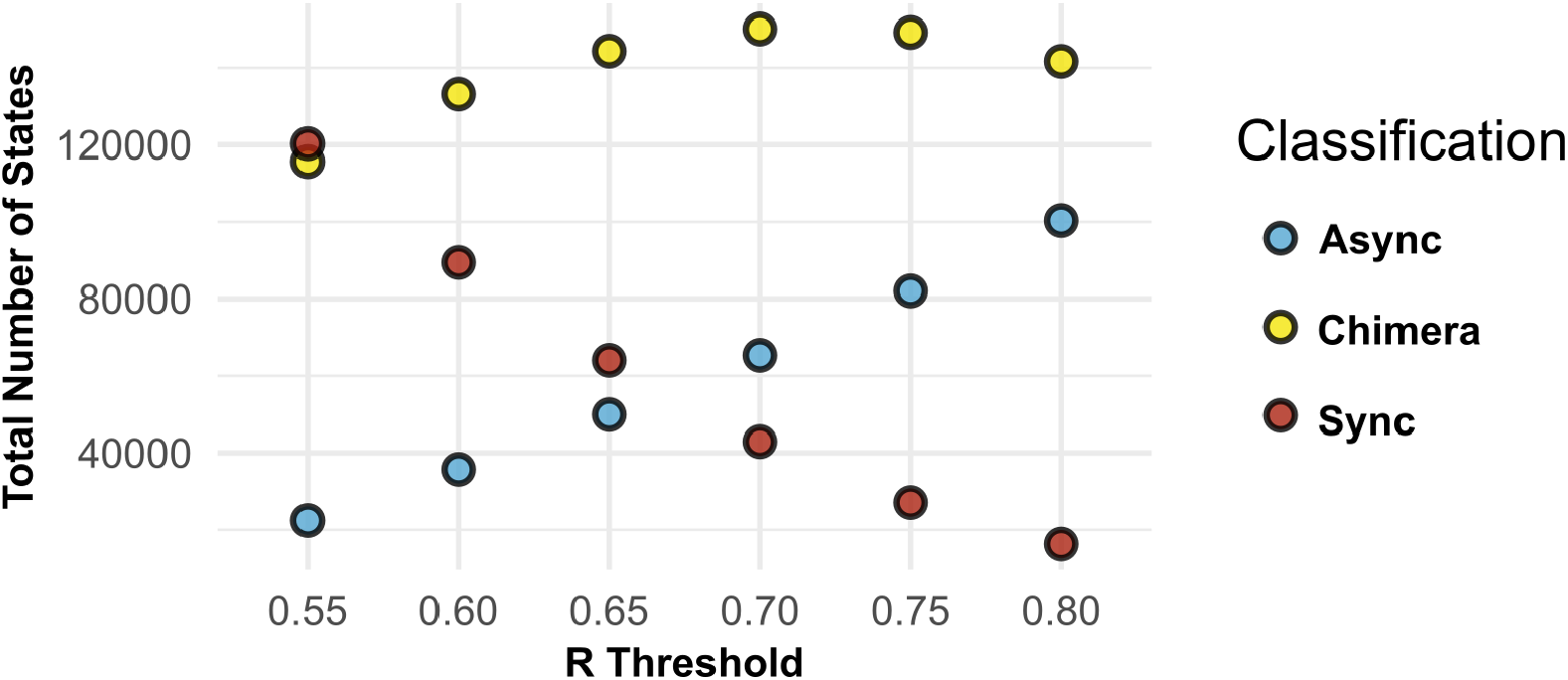
The total number of synchrony classification states produces at different synchrony thresholds for the asynchronous states (blue), chimera states (yellow), and synchronous states (red). We select the threshold at *R* ≥ 0.65 for our analysis.

**Fig S3.**
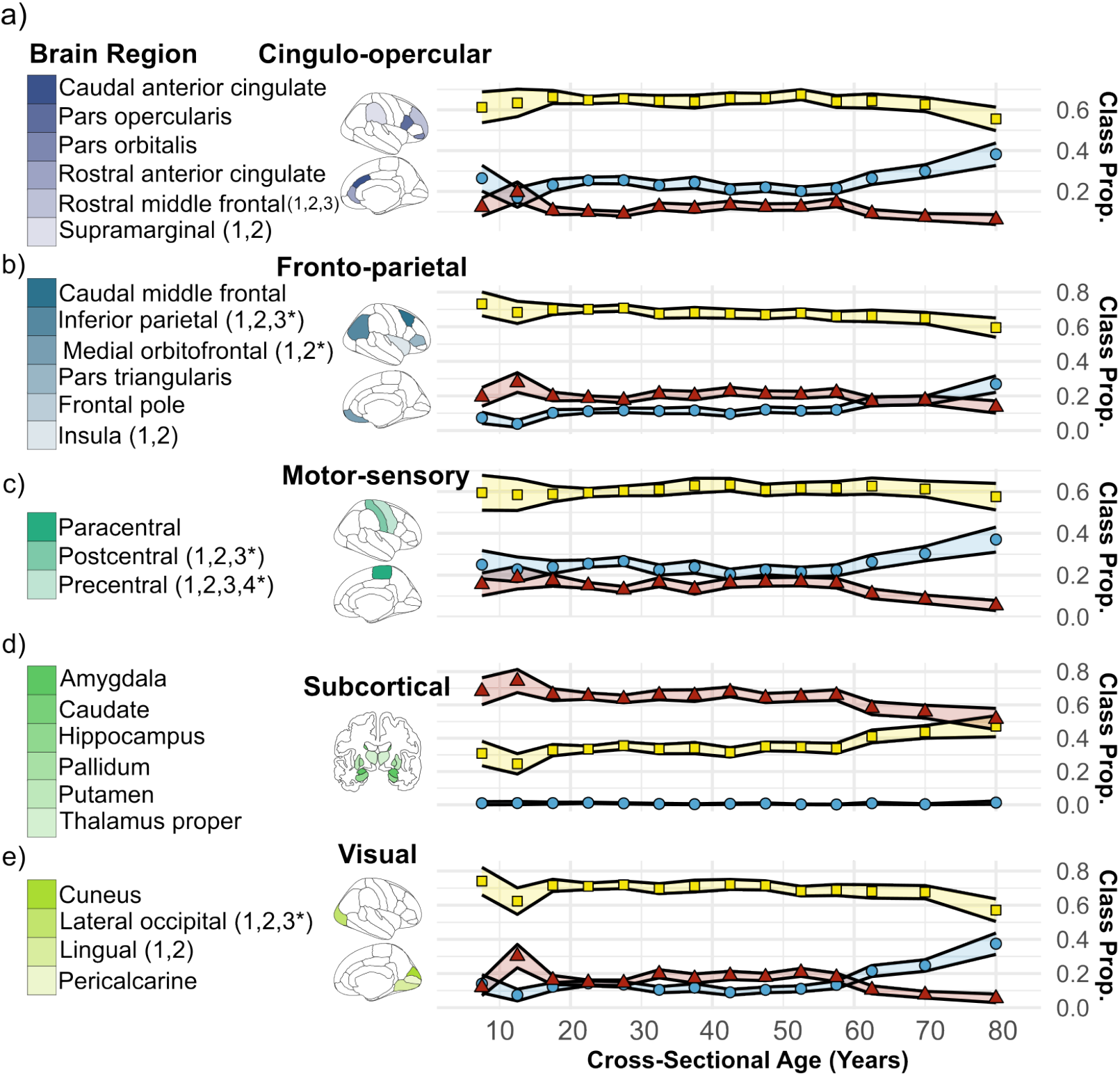
The classification proportions across cross-sectional age for the following cognitive systems: a) cingulo-opercular (blue), b) fronto-parietal (dark blue), c) motor-sensory (teal), d) subcortical (green), and e) visual (light green). Blue circles represent the asynchronous classification proportions, yellow squares represent the chimera state classification proportions, and red triangles represent the synchronous classification proportions. The first column shows the brain region labels that belong to the respective cognitive system for each cognitive system, the second column shows where the brain regions are located spatially, and the third column shows how the classification proportions change with cross-sectional age. Shaded regions are 95% confidence intervals across individuals. *The number of subregions belonging to a brain region may differ depending on the hemisphere. In this case, the left-inferior parietal has 2 and the right-inferior parietal has 3, the left-medial orbitofrontal has 1 and the right-medial orbitofrontal has 2, the left-postcentral has 3 and the right-postcentral has 2, the left-precentral has 4 and the right-precentral has 3, and the left-lateral occipital 2 and the right-lateral occipital has 3.

**Fig S4.**
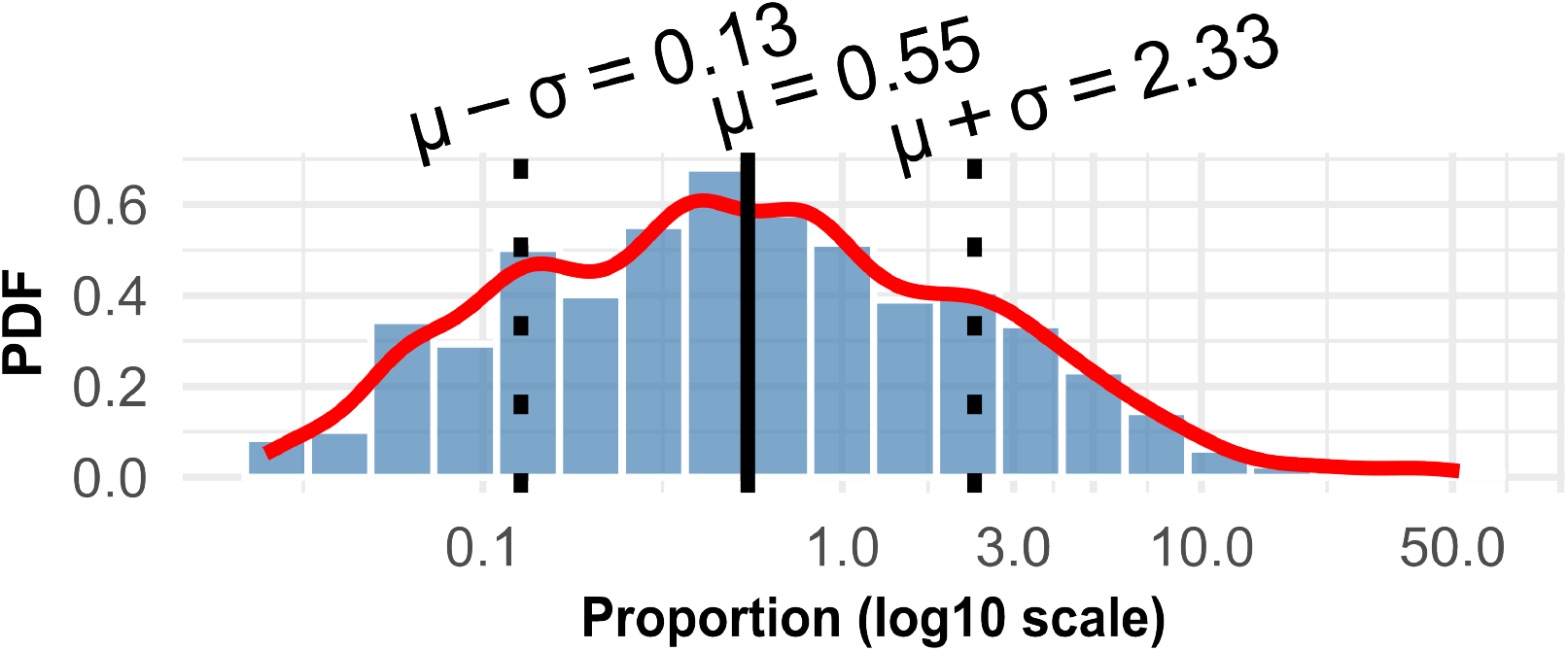
The probability density function of all synchronization pattern proportions in log 10 scale with the mean *μ* (solid black line) and standard deviation *σ* (dotted black line). A kernel density estimator is shown in red.

**Fig S5.**
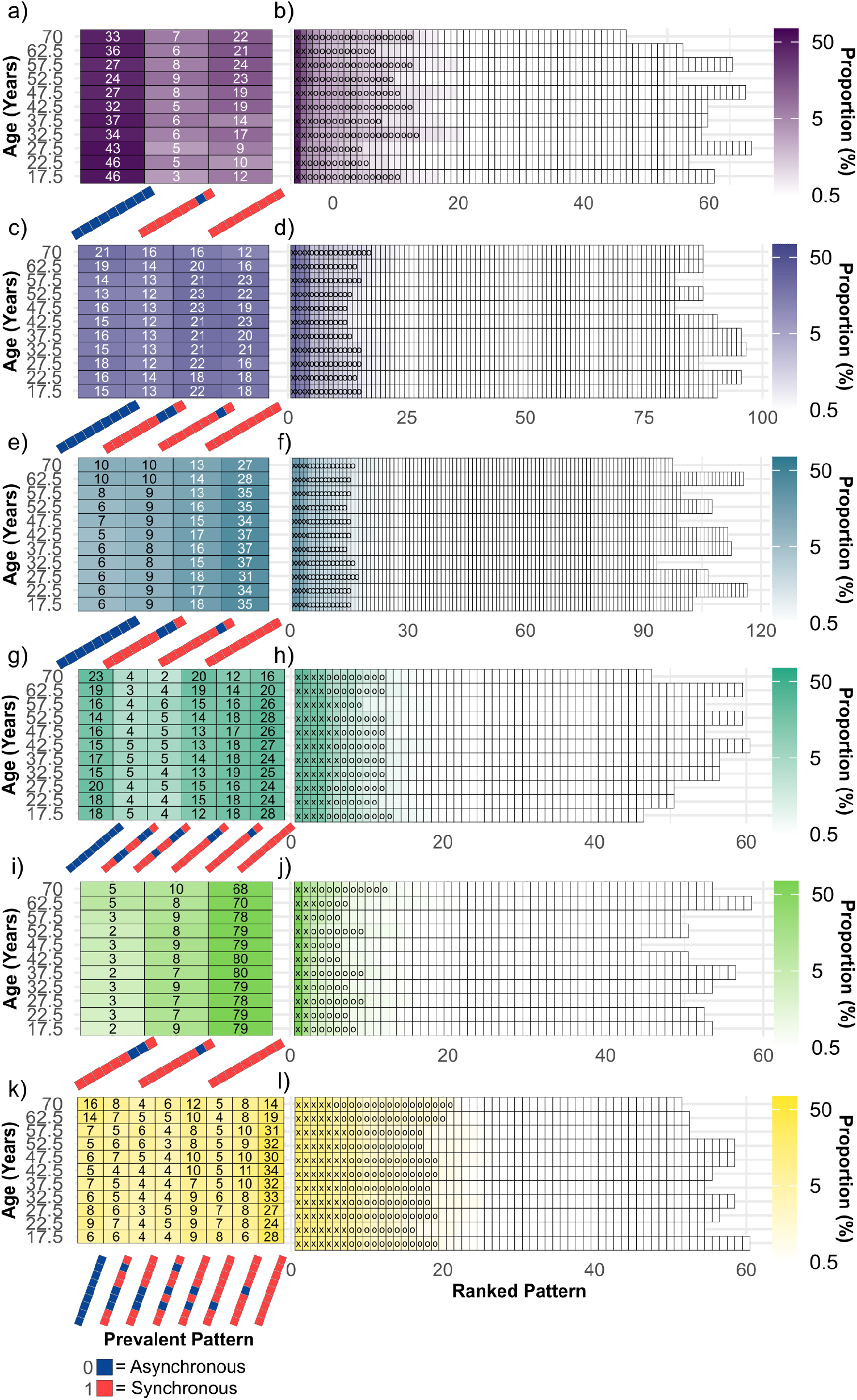
Same as Fig. 8 but for the auditory (panels a-b; dark purple), cingulo-opercular (panels c-d; purple), fronto-parietal (panels e-f; blue-grey), motor-sensory (panels g-h; teal), subcortical (panels i-j; green), and visual (panels k-l; yellow) cognitive systems.

**Fig S6.**
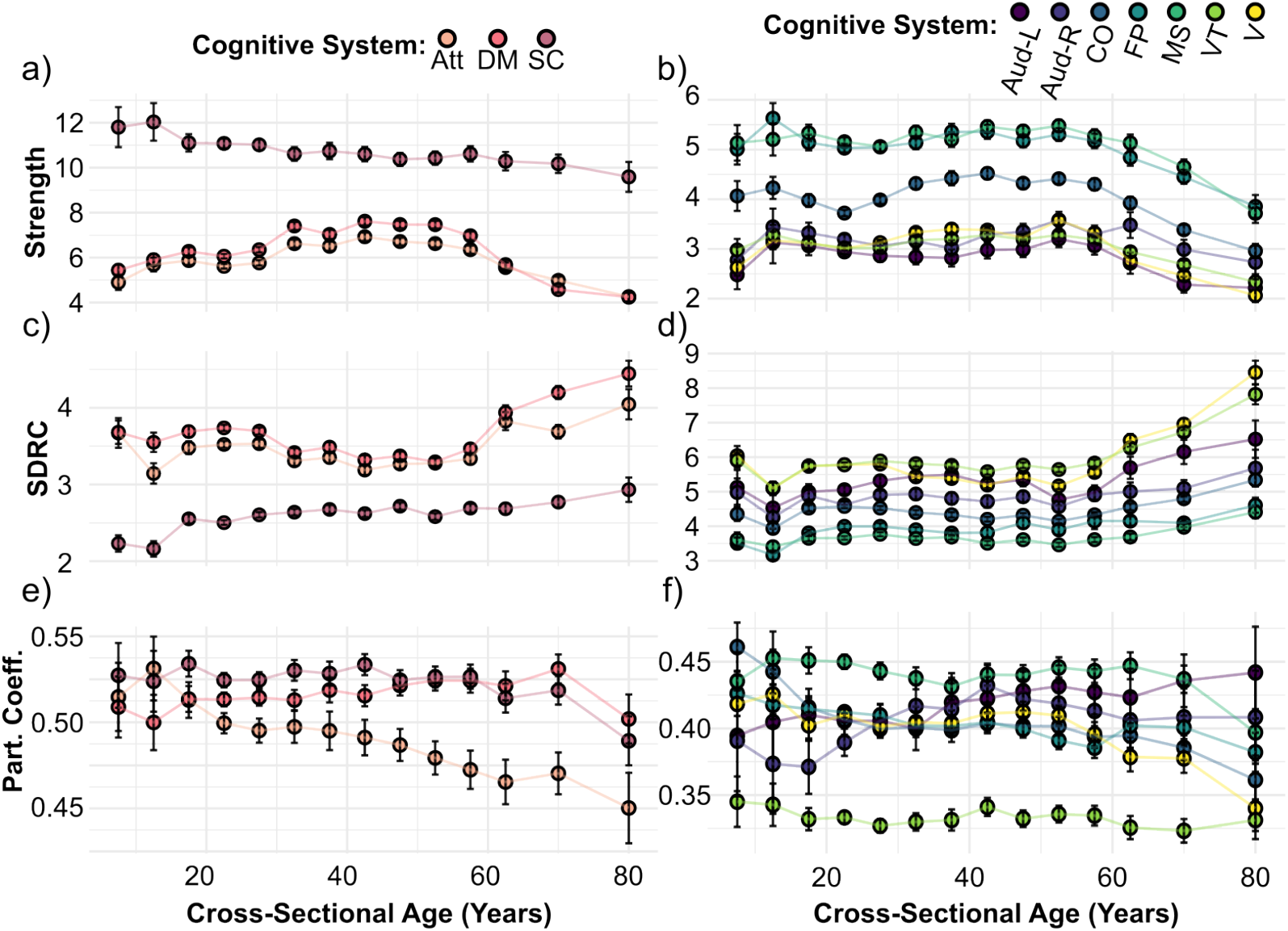
Topological measures of the structural connectomes over cross-sectional age: The average strength (panels a and b), the shortest-distance rich-club (SDRC; panels c and d), and the participation coefficient (panels e and f). Each color represents a different cognitive system, for panels a and c, orange for attention, pink for default mode, and dark magenta for subcortical and for panels b and d, dark purple for left-hemisphere auditory, purple for right-hemisphere auditory, blue-grey for cingulo-opercular, teal for fronto-parietal, dark green for motor-sensory, green for ventral-temporal, and yellow for visual. Error bars correspond to the standard error of the mean at the 2*σ*-level.

**Fig S7.**
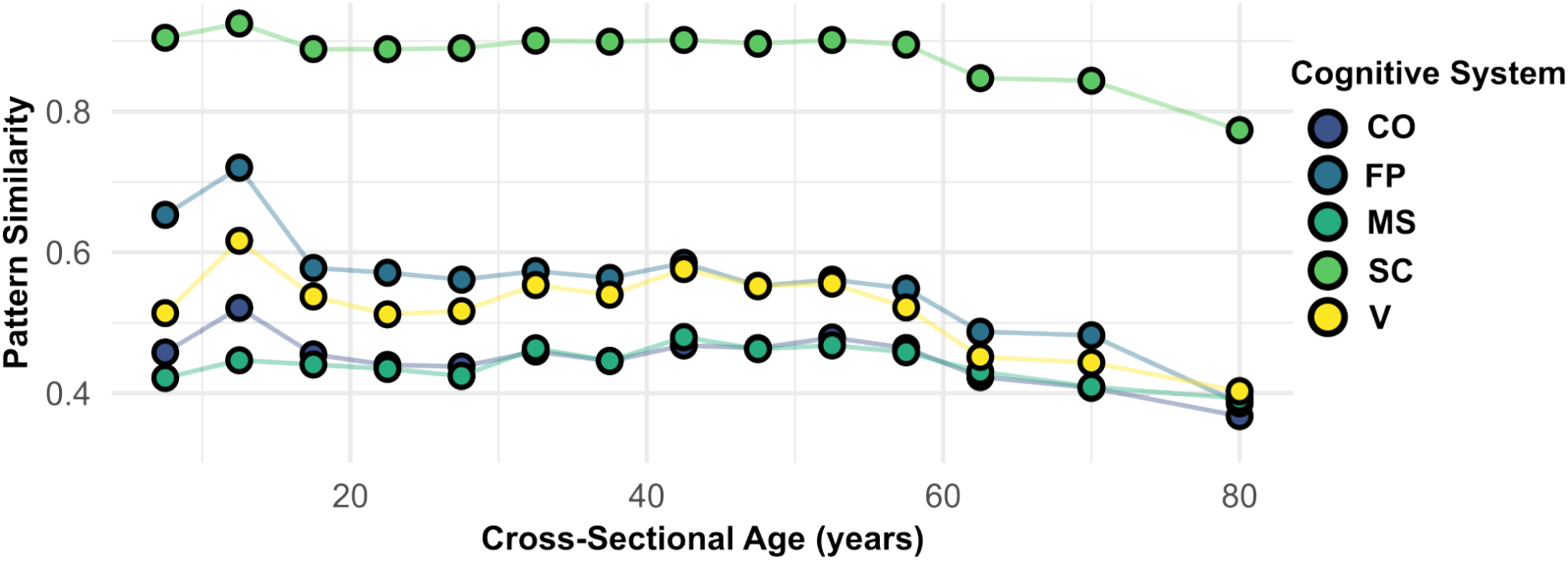
Variations in similarity of synchronization patterns, as defined in Eq. 12, with cross-sectional age across the Cingulo-opercular (CO; blue), Fronto-parietal (FP; dark blue), Motor-sensory (MS; teal), Subcortical (SC; green), and Visual (V; yellow) cognitive systems.

**Fig S8.**
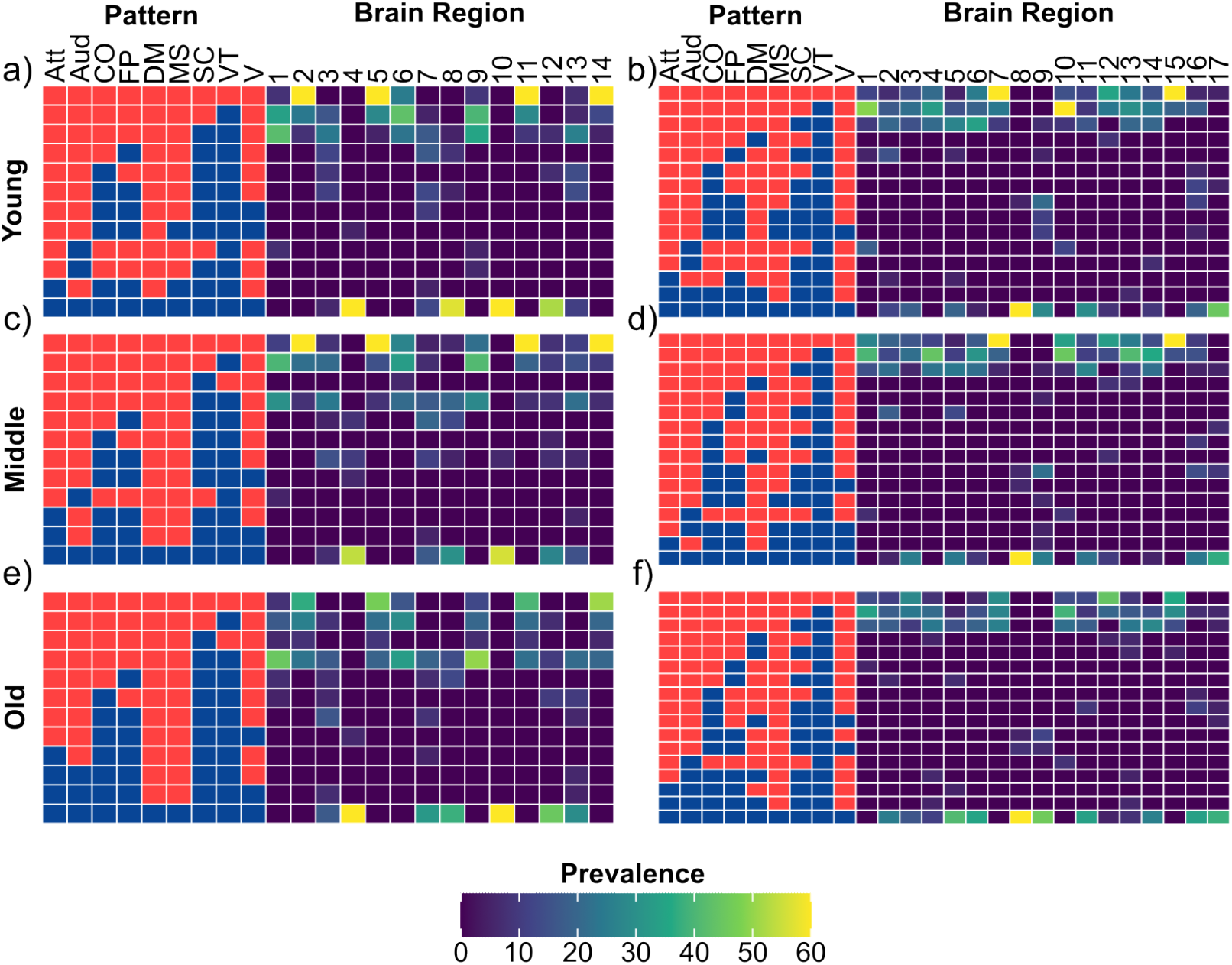
The prevalence of synchronization patterns for each brain region belonging to the motor-sensory cognitive system (the first column, panels a,c,e) and cingulo-opercular cognitive system (the second column, panels b,d,f). The top (panels a,b), middle (panels c,d), and bottom row (panels e,f) represent the age group young (*<* 30 years), middle (≥ 30 & *<* 60 years), and old age (≥ 60 years), respectively. The columns titled with pattern represent the synchronization pattern and are colored red for synchronous and blue for asynchronous grouping with the left-to-right columns representing the attention (Att), auditory (Aud), cingulo-opercular (CO), fronto-parietal (FP), default mode (DM), motor-sensory (MS), subcortical (SC), ventral-temporal (VT), and visual (V) cognitive systems, respectively. The columns titled with brain region each represent a single brain region per column and are colored according to the prevalence of the emergent synchronization pattern when stimulating that brain region across all individuals in a given age group.

**Fig S9.**
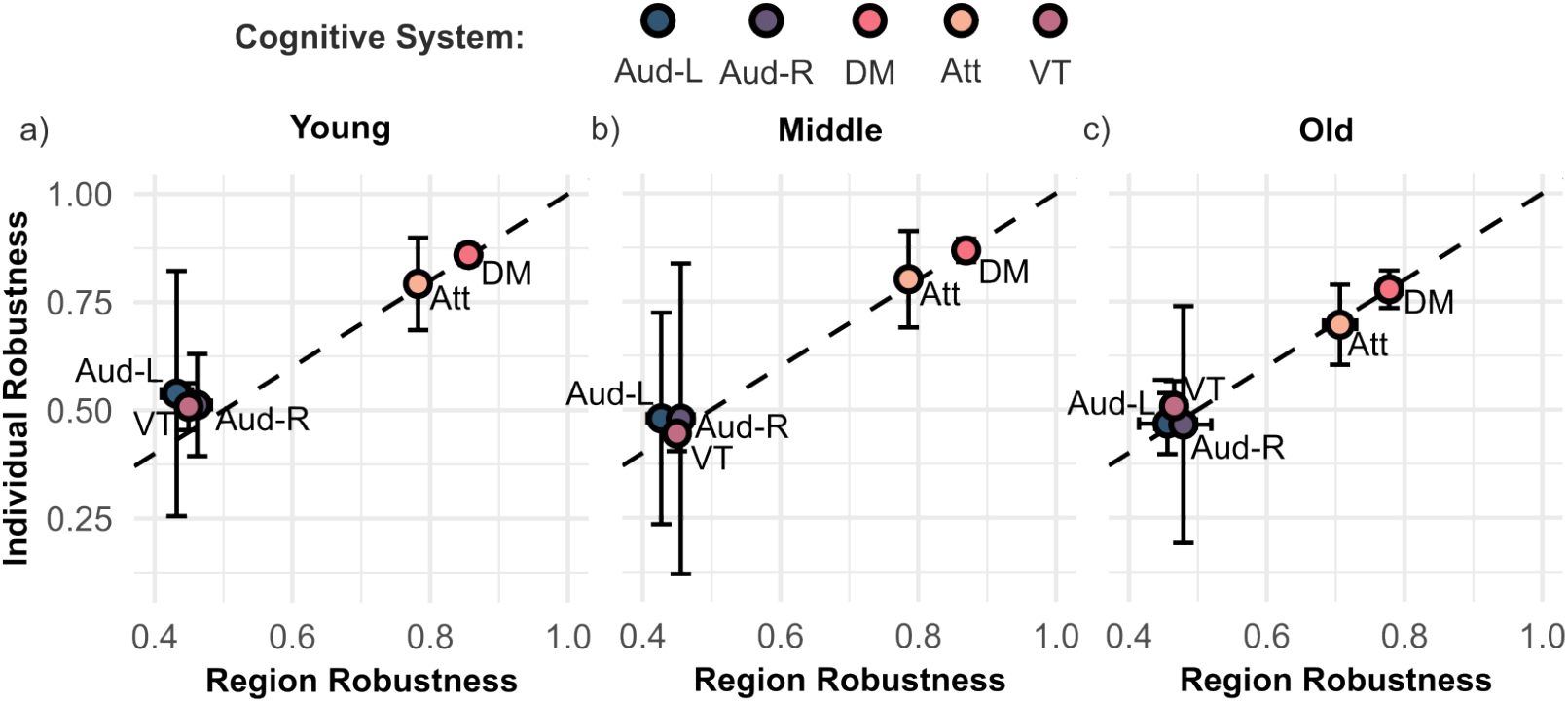
Individual robustness and region robustness across the three age groups: a) young (*<* 30 years, b) middle-aged (≥30 & *<* 60 years), and c) old age (≥60 years) for five cognitive systems: auditory-left (Aud-L; dark blue), auditory-right (Aud-R; dark purple), ventral-temporal (VT; dark magenta), default mode (DM; pink), and attention (Att; orange).

## Footnotes

1 Note that it is possible that there are two or more communities that consist of more than a single member. This, however, only occurs on rare occasions, namely for *≈* 1.7% of the total number of cases.

2 We have directly tested our computational pipeline on the data used in Ref. [52] and largely reproduced their findings.

## Notes

### Competing Interest Statement

The authors have declared no competing interest.

